# Flexible use of conserved motif vocabularies constrains genome access in cell type evolution

**DOI:** 10.1101/2024.09.03.611027

**Authors:** Chew Chai, Jesse Gibson, Pengyang Li, Anusri Pampari, Aman Patel, Anshul Kundaje, Bo Wang

## Abstract

Cell types evolve into a hierarchy with related types grouped into families. How cell type diversification is constrained by the stable separation between families over vast evolutionary times remains unknown. Here, integrating single-nucleus multiomic sequencing and deep learning, we show that hundreds of sequence features (motifs) divide into distinct sets associated with accessible genomes of specific cell type families. This division is conserved across highly divergent, early-branching animals including flatworms and cnidarians. While specific interactions between motifs delineate cell type relationships within families, surprisingly, these interactions are not conserved between species. Consistently, while deep learning models trained on one species can predict accessibility of other species’ sequences, their predictions frequently rely on distinct, but synonymous, motif combinations. We propose that long-term stability of cell type families is maintained through genome access specified by conserved motif sets, or ‘vocabularies’, whereas cell types diversify through flexible use of motifs within each set.

## Introduction

Cell type specialization is a hallmark of multicellular life, allowing diverse functions to co-evolve in the same genome^1,2^. Decades of research on cell types have revealed several fundamental principles. First, cell types are hierarchically organized into families (e.g., neurons) containing evolutionarily related types (e.g., hunger or thirst neurons) with similar gene expression profiles^3,4^. Second, these families remain recognizable across long evolutionary distances based on their conserved gene expression signatures^5,6^, even when species-specific modifications erode one-to-one cell type correspondence between species^1,6^. Third, cell type identities are encoded by the genome. To specify multiple cell types from a single genome, only a part of it is accessible for transcription and regulation in individual cell types^7^. In this way, genomes can be thought of as partially-opened catalogs, from which cells can pick the molecular machinery to assemble themselves.

Synthesizing these principles raises the question of how genome accessibility conveys the stability and hierarchy of cell type identities. Pioneering comparative single-cell epigenomic analysis of the mammalian neocortex showed that ∼50% of the genome regions accessible in human neocortex cells have conserved sequences in other mammalian genomes, but only ∼18% are also accessible in other mammals’ neocortices and merely ∼4% show matched patterns across cell types, despite a divergence of less than 90 million years among the studied animals^8^. This suggests that genome accessibility undergoes turnover even faster than the sequence itself. These observations align with the functional studies showing that point mutations in enhancers can drastically alter the specificity of gene regulation, even without completely disrupting transcription factor (TF) binding^9,10^. Recent work comparing homologous cell types between the mammalian neocortex and chicken pallium identified ‘enhancer codes’ (fixed combinations and arrangements of TF binding sites) within accessible genome regions otherwise lacking sequence conservation^11^. These enhancer codes drove accessibility patterns that were conserved in some cell types, yet lost specificity in others^11^. Given that non-coding sequences quickly diverge^12^ and gene order can be extensively scrambled^13^, it is unclear to what extent genome accessibility may be conserved across longer evolutionary distances and what information the conserved elements convey.

We sought to address these questions using two groups of early-branching animals – flatworms and cnidarians – which have diverged for ∼500-600 million years within each group (**Figure 1A**). We collected full-body multiomic single-cell atlases for three flatworm species: the marine flatworm *Macrostomum lignano*, the freshwater planarian *Schmidtea mediterranea*, and the human parasite *Schistosoma mansoni*, measuring gene expression (RNAseq) and genome accessibility (ATACseq) simultaneously. We showed that most major cell type families shared by these species do retain conserved chromatin accessibility patterns. Using deep learning models, we extracted conserved sets of sequence motifs dictating the accessibility in each family and demonstrated *in-silico* that these motif sets are sufficient in differentiating accessibility between families. Surprisingly, we found that the combinatorial interactions between these motifs defining cell type relationships are species-specific, a result that appears to be reproducible in our comparison of two cnidarians – *Hydra vulgaris* and *Nematostella vectensis* – implying minimal conservation of cell type-specific enhancer codes at these longer distances.

**Figure 1.**
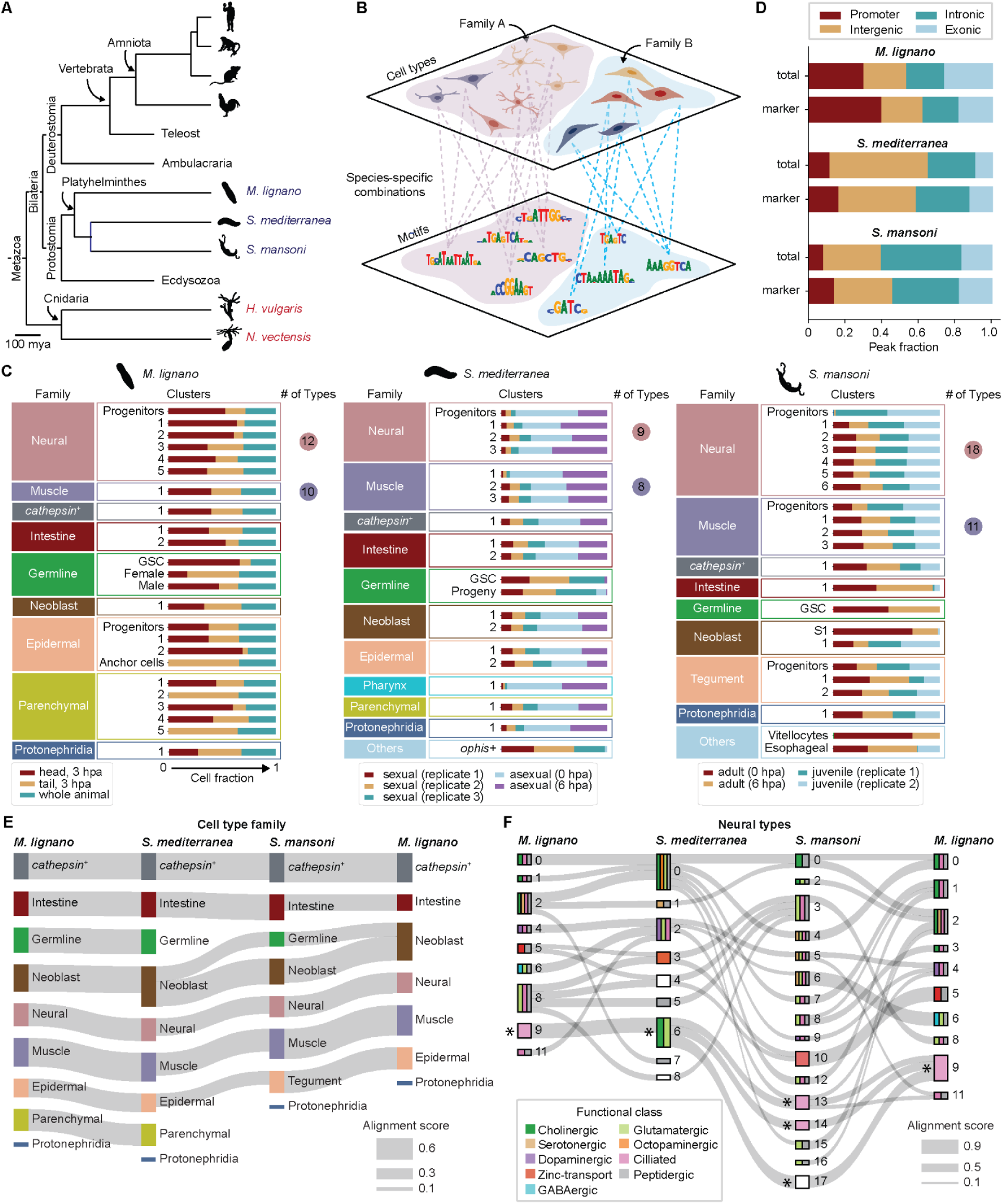
Chromatin accessibility is conserved within cell type families. (A) A schematic, simplified phylogeny including flatworms (blue) and cnidarians (red) studied in this work. Representative amniote species used in previous studies^8,11^ are also included. (B) Schematic of the proposed ‘collective maintenance’ model. Sets of motifs with conserved accessibility biases in broad cell type families (bottom) collectively specify accessibility within those families (top). Species-specific combinatorial motif usage defines fine-grained cell type accessibility patterns. Cells are colored to represent distinct species. (C) Stacked bar plots reporting the proportions of cells within each annotated cluster across sequencing libraries. Clusters for each species are grouped into major cell type families, and the number of neural and muscle types identified through subclustering are indicated to the right of their respective family. While sample compositions are largely uniform, exceptions are observed consistent with expected sampling biases. Tail-specific anchor cells, as well as several parenchymal clusters, which may represent tail-specific glands in *M. lignano*^22^, are not captured in the head dataset. The germline stem cells, their progeny, and accessory cells (*ophis+*, S1, vitellocytes)^15,21^ are not captured in the asexual planarian data, nor the sexually immature juvenile schistosome data. Sexual planarian samples are depleted in many somatic clusters as these were dissected to enrich for the reproductive organs. hpa: hour post amputation. (D) Fractions of all accessible peaks (top) and cluster marker peaks (bottom, **Figure S2A**) that are either proximal (promoter, exonic, intronic), or distal (intergenic) to gene bodies, showing no enrichment of any category in marker peaks. (E) Sankey plot summarizing the chromatin accessibility-based mappings between flatworm cell type families using SAMap. Edges with alignment score <0.3 are omitted. *M. lignano* cell types are shown on both sides to show all pairwise comparisons. Functionally conserved families such as *cathepsin^+^* phagocytes, intestinal cells, muscles, neurons, and neoblasts clearly map between species. Planarian and *Macrostomum* epidermal cells map to schistosome tegument, which is the parasite’s syncytial epidermis^15^. Planarian parenchymal cells map to previously reported gland cells^22^ in *Macrostomum*, such as the prostate and cement glands (**Figure S1A** and **S2C**), but appear to be lost in the schistosome. (F) Sankey plot showing highly degenerate mappings between flatworm neural cell types. Edges with alignment score <0.2 are omitted. Blocks representing cell types are colored to indicate their annotated functional classes (**Figure S1B**). Populations containing neural progenitors are marked with an *, and map strongly to one another despite expressing genes associated with distinct functional identities.

Our results support a ‘collective maintenance’ model (**Figure 1B**): sequences drawing from separate, conserved sets of motifs – or vocabulary in linguistic terms – convey family-level accessibility biases independent of specific motif choice and combination. This flexibility may confer long-term stability to family identities, while enabling diversification of new cell type-specific accessibility patterns through combinatorial motif use.

## Results

### Cell type families show conserved chromatin accessibility landscapes

We collected single-nucleus multiomic data from whole-body samples of the three flatworms, retaining ∼90,000 nuclei after strict quality control (see **Methods**). This dataset encompassed various biological conditions for each species, including different reproductive biotypes, developmental stages, and injury states (**Table S1**), allowing us to focus on cell type-defining features independent of these factors. Integrating ATACseq and RNAseq data, we identified major cell clusters spanning 8 conserved cell type families shared by all three species, as well as the parenchymal glandular cell type family lost in *S. mansoni*, based on known marker genes^14–19^ (**Figures 1C** and **S1**). We also captured species-specific rare cell types, including *foxA^+^* schistosome esophageal gland cells^20^, *ophis^+^* planarian germline accessory cells^21^, and *macif1^+^* anchor cells^22^ from the *Macrostomum* adhesive organ. Together, our dataset provides a broad coverage of cell type diversity within these animals.

Comparing chromatin accessibility patterns between these species is challenging due to substantial sequence divergence and structural variations^23^. Therefore, we tested whether the local ATACseq signal around homologous gene bodies (i.e., gene scores) is comparable. This approach was feasible because we observed that ATACseq peaks specific to each cell cluster (**Figure S2A**) contained similar proportions of promoters, exonic, intronic, and intergenic peaks as the complete peak sets (**Figure 1D**). Logistic regression models trained to annotate cell clusters based on peak accessibilities also revealed no significant differences in the predictive power of these peak types (**Figure S2B**). These observations suggest that regulatory information associated with cell type identity is present in regions both proximal to and distal from gene bodies.

**Figure 1E** shows that most shared cell type families can be linked uniquely across species based on accessibility using gene scores through SAMap^6^, a tool designed to identify cell type homologies based on shared gene expression patterns. Consistently, species-specific clusters stayed unconnected (**Figure S2C**). Protonephridial cells and schistosome germline cells were notable exceptions – showing negligible mappings or cross-family mappings, respectively, despite conserved function and marker gene expression^15,19,24,25^.

To examine whether this conservation extended to cell types, we sub-clustered neurons for each species, identifying populations with cholinergic (*chat*^+^/*vacht*^+^), serotonergic (*sert*^+^), dopaminergic (*th*^+^/*ddc*^+^), and peptidergic (*pc2*^+^/*7b2*^+^) signatures among others^26^ (**Figure S1B**). In addition, we captured *znt2*^+^/*zip6*^+^ neurons in all three species, corresponding to previously annotated planarian *otf*^+^ neurons^17^ and schistosome *kk7*^+^ neurons^15^, which we termed zinc-transport neurons. Comparing these neural types between species, SAMap reported highly degenerate mappings among differentiated neurons (**Figure 1F**), though progenitors were almost exclusively mapped to one another (**Figure 1F**). We also compared muscle types between species using the same approach and observed similar multi-mappings (**Figure S2D**). These results contrast the one-to-one cross-species mappings at the family level, suggesting that the chromatin accessibility is most strongly conserved within cell type families rather than between individual cell types.

### Deep learning models identify conserved motif usage in accessible genomes

To reveal the sequence features driving these chromatin accessibility patterns, we used ChromBPNet models^27,28^. These models predict ATACseq insertion counts at base pair resolution from the underlying sequences, after regressing out noise induced by sequence preferences of the Tn5 enzyme, which vary with GC content (**Figure S3A**). Using model interpretation tools (see **Methods**), we identified sequences driving predictions of the residual accessibility, which often correspond to TF binding sites^27^. Unlike conventional motif matching analysis, this approach is not constrained by a predefined set of motifs. Instead, the models identify these sequences de novo and filter out potential TF binding sites that are unlikely to contribute to the observed accessibility^29^ – an important feature for non-model species lacking abundant ChIPseq data.

To illustrate our analysis, we examined the loci of *nanos* homologs – deeply conserved RNA binding proteins involved in the germline development of diverse animals^30^. Our previous work showed that *S. mansoni* has two *nanos* homologs, with *Sma-nanos-1* specifically expressed in germline stem cells (GSCs) and required for their maintenance^31^. We trained a ChromBPNet model on aggregate GSC ATACseq data and used DeepSHAP^32^ to attribute the model’s predictions to individual bases in two GSC-specific peaks near the *Sma-nanos-1* locus (**Figure 2A**). Within the first peak, we observed a pair of sequences with high contribution scores (i.e., SHAP scores), matching the binding motif of the NFY TF family, consistent with previous work showing that knockdown of *Sma-nfy-1* ablates the schistosome male germline^33^. Interpreting the second peak revealed a triplet of binding sequences for the JUN/FOS family of basic leucine zipper (bZIP) TFs. Genome-wide analysis showed enrichment of this motif in accessible chromatin regions of the schistosome GSCs, though similar enrichment was not seen in the other two species (**Figure S3B**). Testing the functional relevance of these observations, we performed an RNAi screen of all 13 bZIP domain-containing genes (**Table S2**) in the *S. mansoni* genome and identified one – *Smp_335650 –* for which knockdown specifically ablated the male germline while leaving neoblasts, the flatworm somatic stem cells, unaffected (**Figure 2B**), phenocopying *Sma-nanos-1* RNAi^31^. Similarly, our analyses of *Sma-nanos-2* suggested that *p53* may bind to a regulatory sequence associated with this locus specifically in neoblasts (**Figure 2C**), consistent with previous functional studies^31,34^. These results demonstrate that the trained ChromBPNet models can identify biologically meaningful features in accessible genome regions.

**Figure 2.**
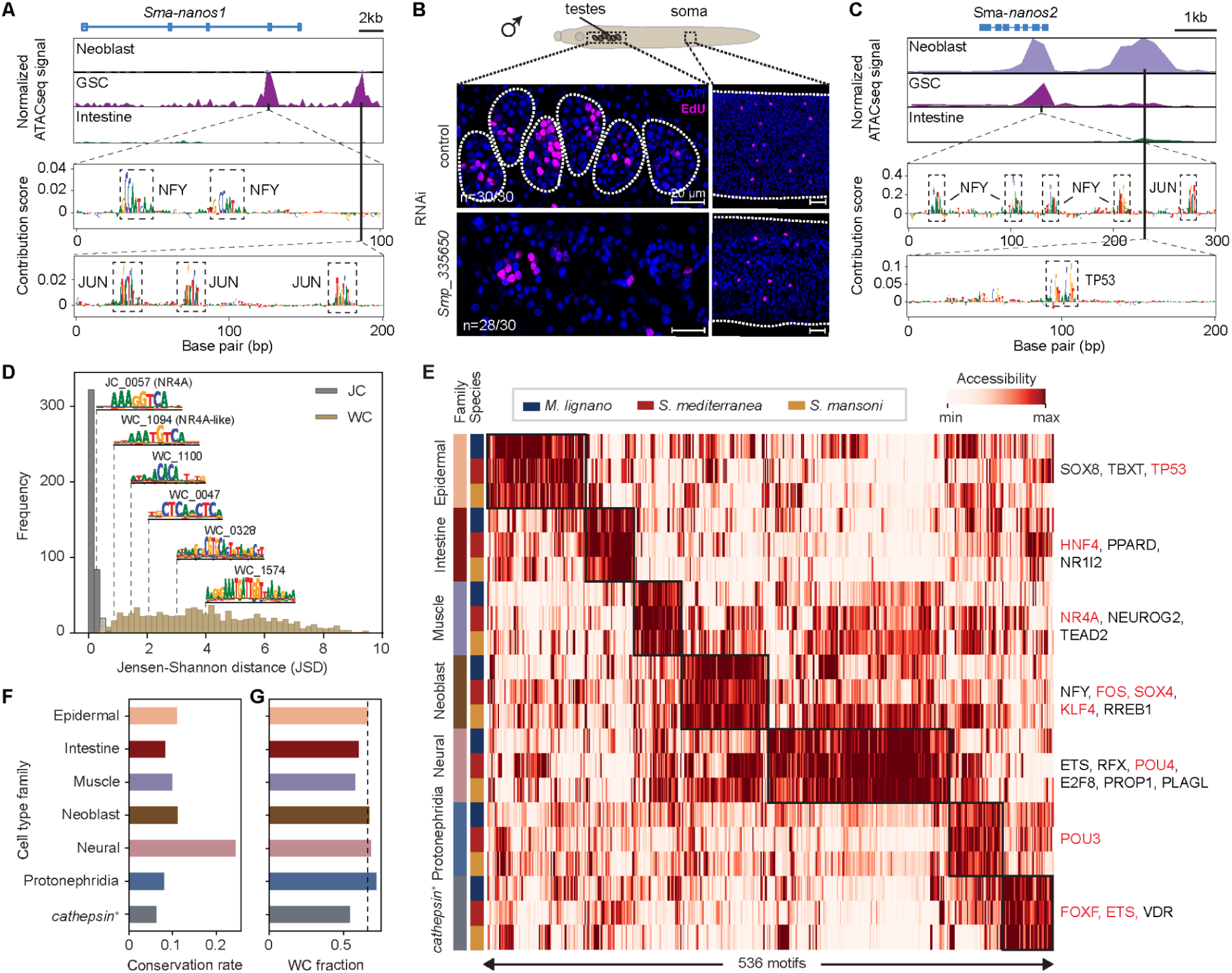
ChromBPNet models reveal sequence motifs with conserved cell type family biases. (A) Top: genome tracks showing pseudo-bulk ATACseq profiles around *Sma-nanos-1* locus (on the reverse strand) normalized such that all groups have the same total counts within the transcription start site (TSS) region, comparing coverage in neoblasts, GSCs, and differentiated cells (using intestine as an example). ATACseq peaks are detected specifically in GSCs, consistent with the GSC-specific expression of *Sma-nanos-1*^19^. Bottom: per-base contribution scores of the peak regions showing short sequences with high importance, matching NFY (top) and JUN (bottom) binding motifs, respectively. (B) Confocal images of testes (left) and somatic tissues (right) in schistosome male juveniles, stained with DAPI (blue) and EdU (magenta) in control and *Smp_335650* RNAi animals. Dashed boxes in the schematic correspond to the imaged areas. Dashed circles in the left images highlight the testis globules, which are absent in *Smp_335650* RNAi animals. Dashed lines in the right images indicate the animal surface. Numbers represent counts of animals that had phenotypes consistent with the images shown out of all animals examined. (C) Top: genome tracks around *Sma-nanos-2* locus (on the reverse strand). While a common peak is present in both neoblasts and GSCs, an additional peak is specifically accessible in neoblasts upstream of the gene, consistent with the expression of *Sma-nanos-2* in both neoblasts and GSCs^31^. Bottom: per-base contribution scores of the peak regions showing sequences matching NFY and JUN motifs in the peak shared by neoblasts and GSCs (top), and a TP53 binding motif in the neoblast-specific peak. (D) Histogram of the Jensen-Shannon distance (JSD) between consensus motif set used in this study and their respective closest matches in the JASPAR2020 database, with representative logos for motifs examined using *in-silico* footprinting (**Figures S3E-F**). Note that the top two motifs only differ by one nucleotide, but show distinct footprint patterns. (E) Min-max normalized average accessibility of 536 motifs (columns) in 7 cell type families shared by all three species (rows), showing conserved family-level accessibility biases (boxes). Example motifs, including those corresponding to known fate specific TFs (red), are listed to the right. Germline is excluded due to lack of conserved marker motifs. (F) Motif conservation rate per-family, calculated as the fraction of motifs that are family markers for all three species, divided by the total number of marker motifs in any species. (G) Fractions of conserved motifs that are WC motifs in each cell type family. Dashed line indicates the fraction of WC motifs in the complete motif set.

We trained ∼30 species-specific models on all major cell type families, combining progenitors with their respective differentiated cell types. We also grouped rare cell types to achieve coverage depth needed for training (see **Methods**). Performance was consistent across three train-test splits, and the resulting models had quality on-par with previous analyses in model organisms^29,35^ (**Figure S3C**). Using TF-MoDISco^36^, we extracted sequence motifs dictating genome access learned by the model and combined them with conserved TF-binding motifs from the reference database JASPAR^37^ to create a minimally redundant set containing 429 JASPAR motifs (labeled as JC motifs) and 840 learned motifs (labeled as WC motifs) (**Figure 2D**, **File S1**). Using *in-silico* footprinting^28,29^, we confirmed that our models are highly sensitive, and could differentiate sequences with single nucleotide differences (**Figures S3D-E**). The models also predicted footprints for some WC motifs, suggesting that these may represent protein binding sequences (**Figure S3F**). To determine whether WC motifs were informative in distinguishing cell types, we trained logistic regression models to annotate major cell clusters based on motif deviations, and found no differences in the predictive power of JC and WC motifs (**Figure S3G**). These results demonstrate that our analysis identified sequence features associated with cell type identities beyond known TF-binding motifs.

To test whether motif usage is conserved across species, we identified ‘marker motifs’ of each major cell cluster using chromVAR deviations^38^ (see **Methods**), and compared them across species. We found ∼600 motifs with conserved biases towards specific cell type families (**Figure 2E, Table S3**). Neurons, together with their progenitors, showed the most conserved motif use (**Figure 2F**), with nearly 25% of marker motifs shared between species. The conserved marker motifs contained both WC and JC motifs (**Figure 2G**), suggesting that they play comparable roles in driving genome accessibility. Many of these motifs corresponded to fate-specifying TFs, such as *p53* in the epidermal lineage^34^, *hnf4* in the intestine^16^, and *foxF* in the *cathepsin^+^* cells^39^ (**Figure 2E**), which have been experimentally characterized in at least one of these animals. This lends strong support to our results, and suggests that the functions of these TFs in establishing cell type identities may be broadly conserved.

### Neurons, muscles, and glandular cells form a cell type super-family based on conserved motif usage

To further characterize the conservation of motif use without presuming cell type relationships, we quantified similarities between all major cell clusters by calculating correlations of average motif deviations between each pair across all species. Clustering this similarity matrix, we found that even when mixing species, members of the same cell type families were generally more similar to one another than to other families (**Figure 3A**). This observation reaffirmed that motif usage is highly conserved at the family level, although species biases prevailed when comparing finer clusters. Strikingly, this analysis revealed a prominent division between two super-families, separating neurons, muscles, and parenchymal cells from all others (**Figure 3A**). Consistent with this division, we found that, while *S. mansoni* lost parenchymal cell types (**Figures 1E**), the conserved marker motifs of *M. lignano* and *S. mediterranea* parenchymal cells were predominantly accessible in the neurons and muscles of *S. mansoni* (**Figure S4A, Table S3**).

**Figure 3.**
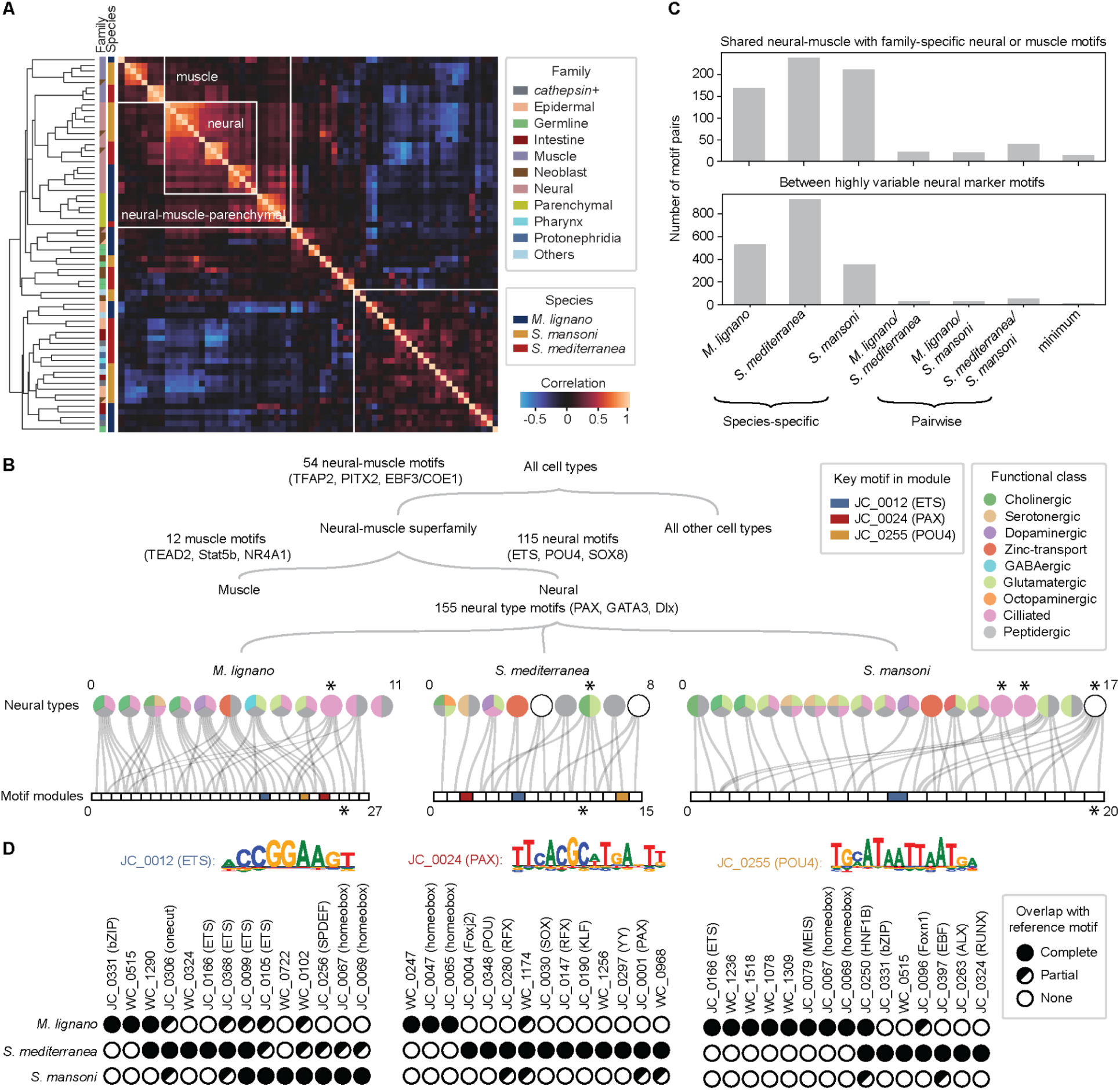
Motif usage defines hierarchy among cell types but combinatorial motif use is barely conserved. (A) Hierarchically clustered pairwise correlations between z-scored average motif deviation profiles for all major cell clusters across three flatworms. Inner color bar: species; outer color bar: cell type family. A superfamily containing neurons, muscles, and parenchymal cells are segregated (upper left box) by strong negative correlations with other differentiated cell types (bottom right box). The germline cell types are at the center, excluded from both super-families. (B) Hierarchical relationships between cell types defined by their motif use, with key motifs labeled on corresponding branches. Neurons and muscles share a conserved enrichment of 54 motifs in their accessible genomes compared to other cells outside of their super family. Within this group, enrichment of 12 motifs distinguish muscles from neurons in all three species, and another 115 are enriched in neurons compared to muscles. Finally, a pool of 155 highly variable marker motifs define species-specific neural types represented by circles and colored by their functional annotations, with progenitors highlighted by asterisks (**Figure S1B**). These motifs associate in species-specific modules with unique accessibility patterns. Non-trivial modules (those containing more than one motif) are represented for each species as a rectangular array with each linked to neural types in which they are accessible (**Figure S4E**), and the major progenitor-specific modules are marked with asterisks. Modules containing ETS (blue), PAX (red), and POU4 (orange) motifs are highlighted with colored boxes in the array. In *S. mansoni*, PAX and POU4 are members of singleton modules not shown here, with high accessibility in neural types 4 & 7, and 7 & 8, respectively. (C) Barplot summarizing number of motif pairs with high co-occurrence enrichment (>4) between neural and muscle shared motifs, and their respective family-specific motifs (top). Species-specific enrichments are drastically reduced in minimum enrichments between each species pair, and almost fully eliminated in the minimum among all three species. Bottom panel shows corresponding values for co-occurrence enrichment between highly variable neural marker motifs. (D) Representations of the modules associated with ETS, POU4, and PAX motifs in each species. Filled circles: identical binary accessibility patterns with the reference motif (same module); half-filled circles: partial overlap, with accessibility differing in one or more neural populations, including both gained or lost accessibility; open circles: non-overlapping binary accessibility.

In support of this neural-muscle grouping, we identified 54 motifs with high deviations in both muscles and neurons for all three species, including TFAP2 (JC_0283), PITX2 (JC_0071), and EBF3/COE1 (JC_0397) (**Figure 3B, Table S3**). Beyond individual motifs, ChromBPNet models trained on muscle and parenchymal data tended to interpret the sequences of neural marker peaks more similarly to neural models than those trained on other cell type families within each species (**Figure S4B**). Similar trends were observed for muscle marker peaks, with neural models producing the most concordant interpretations. Given that neurons and muscles have distinct developmental lineages in flatworms^40^, the observed similarities support the long-standing hypothesis that these families may have a common evolutionary origin^41^, demonstrating a conserved regulatory language alongside their shared molecular machinery.

### Combinatorial motif usage is minimally conserved

We asked how motif usage shapes the relationships between neurons and muscles. For each species, the shared neural-muscle motifs appeared not only in peaks accessible within both families but also in family-specific peaks (**Figure S4C**), suggesting that interactions with other motifs may be necessary to determine their roles in each family. Accordingly, we identified 12 motifs (e.g., NR4A1, Stat5b, TEAD2) with conserved high accessibility in muscles compared to neurons, and 115 motifs (e.g., ETS, POU4, SOX8) with conserved high accessibility in neurons as compared to muscles (**Figure 3B, Table S3**), many of which are consistent with previously characterized muscle/neural regulators^42–44^. These motifs had strong interactions with the shared motifs for each species, measured by co-occurrence^27,45^ enrichment – the ratio between the observed and expected numbers of peaks in which a pair of motifs are annotated together (**Figures 3C** and **S4D**). However, comparing between species, we were surprised to find that a negligible number of these interactions were conserved (**Figure 3C**). In other words, the strongest interactions between pairs of motifs defining the relationships between neurons and muscles were mostly divergent between species.

We wondered whether stricter rules of motif interaction might be necessary to define the relationships between individual cell types in a functionally diverse family like neurons. Comparing motif accessibility among neurons specifically, we identified a set of 155 motifs which were good markers for at least one neural type in each species (**Figure 3B, Table S3**). We defined binarized accessibility profiles for each motif across types (see **Methods**) and grouped them into co-accessible modules (**Figures 3B** and **S4E, Table S3**). Once again, we observed minimal conserved structure in these modules between species. For instance, we noticed that an ETS-family motif (JC_0012) had high accessibility in the zinc-transport neurons in all three species, which all expressed an orthologous ETS-family TF (*etv*) (**Figures S4E-F** and **S5A**). Fourteen motifs had the same binarized accessibility pattern as JC_0012 in at least one species, but none of these were in the same module for all three species (**Figure 3D**). When examining the modules associated with another two highly variable motifs – PAX (JC_0024) and POU4 (JC_0255) – we reached similar conclusions (**Figure 3D**), indicating high flexibility in motif-motif interactions among neural types.

The variability in combinatorial motif use extended to the relationships between TFs and functional effectors. For example, the expression of *pax* homologs (**Figure S5B**) and PAX motif accessibility were associated with GABAergic, dopaminergic, and serotonergic neurons of *M. lignano*, *S. mediterranea*, and *S. mansoni*, respectively (**Figures 3B** and **S4E-F**). Similarly, while *pou4* homologs (**Figure S5C**) were expressed in a subset of peptidergic neurons for all three species, some *pou4*^+^ cells were also glutamatergic in *M. lignano* and *S. mansoni* (**Figures 3B, S4E-F**).

Previous studies observed lower conservation of cis-regulatory elements used in differentiated neurons among mammals compared to progenitor cell types^46^. This led us to examine neural progenitor populations, which were generally linked to higher numbers of motif modules compared to fully differentiated types (**Figure 3B**). The majority of these progenitor motifs, however, belonged to a single, progenitor-specific module for each species (**Figure 3B**), but the overlap between these modules was minimal, with only 3 out of a total of 88 motifs being shared across species (**Table S3**). These results suggest that the progenitors are no more conserved than differentiated cell types in terms of motif use.

Finally, we computed co-occurrence between pairs of neural type-specific motifs, and observed a similar trend of strong species-specific interactions being lost when comparing between species (**Figures 3C** and **S4G**). Collectively, these results support the idea that, although similar sets of motifs are used to define relationships between and within cell type families, the rules for how those motifs combinatorially shape chromatin accessibility are not fixed.

### Divergent combinatorial motif usage is general

To assess the broader relevance of our observations, we expanded our analysis beyond flatworms to cnidarians, which diverged from bilaterians ∼700 million years ago. We analyzed previously published bulk ATACseq data for both neural and non-neural cells collected from *Hydra vulgaris*^47^ and *Nematostella vectensis*^48^ (separated by over 600 million years) using the same pipeline (**Figures S5D-F**). This allowed us to identify 25 motifs with enriched accessibility in the neurons of both cnidarian species, five of which – including POU4 (JC_0255), TFAP2 (JC_0283), PLAG1 (JC_0062), and BACH1 (JC_0193) – were also shared with flatworm neurons (**Figure S5E, Table S3**). These numbers likely underestimate the extent of conservation, as our analysis is limited by low coverage in the *N. vectensis* data. We computed co-occurrence enrichment for the 25 cnidarian neural motifs and observed, similar to flatworms, that interactions were largely specific to individual species (**Figure S5F**). A notable exception was the strong interaction observed between POU4 and Dlx2 (JC_0047), which appeared conserved among the cnidarians but not flatworms. Nevertheless, these results suggest that the principle of conserved motifs interacting under species-specific rules to drive chromatin accessibility applies to more than just flatworms.

Observing this consistent trend of divergent combinatorial motif relationships between species, we asked whether the phenomenon holds at individual loci. We wanted to focus on deeply conserved master regulators, where motif usage may be more tightly constrained, and the prominence of POU4 in our analysis made it a promising candidate. POU4 was preferentially used by neurons of both cnidarians and flatworms, and the gene *pou4* (*BRN3* in vertebrates) has also been shown to specifically express and play essential roles in the nervous system of *N. vectensis*^49^, *S. mediterranea*^40^, *C. elegans*^43^, protovertebrates^50^, and even mice^43^. To compare the regulatory programs around *pou4*, we identified flatworm and *Hydra pou4* homologs (**Figure S5C**), which exhibited well-preserved gene structures – each containing a shorter 5’ exon followed by a longer 3’ exon (**Figure 4A**). Due to low coverage at the *pou4* locus, *Nematostella* was excluded from this analysis. Plotting the ATACseq coverage around these genes, we consistently observed two neural-specific peaks, upstream and downstream of the gene body. Despite these similarities, the motif hits within these regions – ∼60% of which showed a conserved use associated with neural identity – had diverged across species in terms of specific motif choice and arrangements, as well as their relative contributions in defining accessibility (**Figure 4B, File S2**). This was true even of the two planarian paralogs even though both were expressed in neurons. These findings, along with previous documented examples in other animals^51^, demonstrate that regulatory programs associated with deeply conserved transcriptional regulators can also undergo significant modifications while maintaining accessibility in the same cell type family across long evolutionary times.

**Figure 4.**
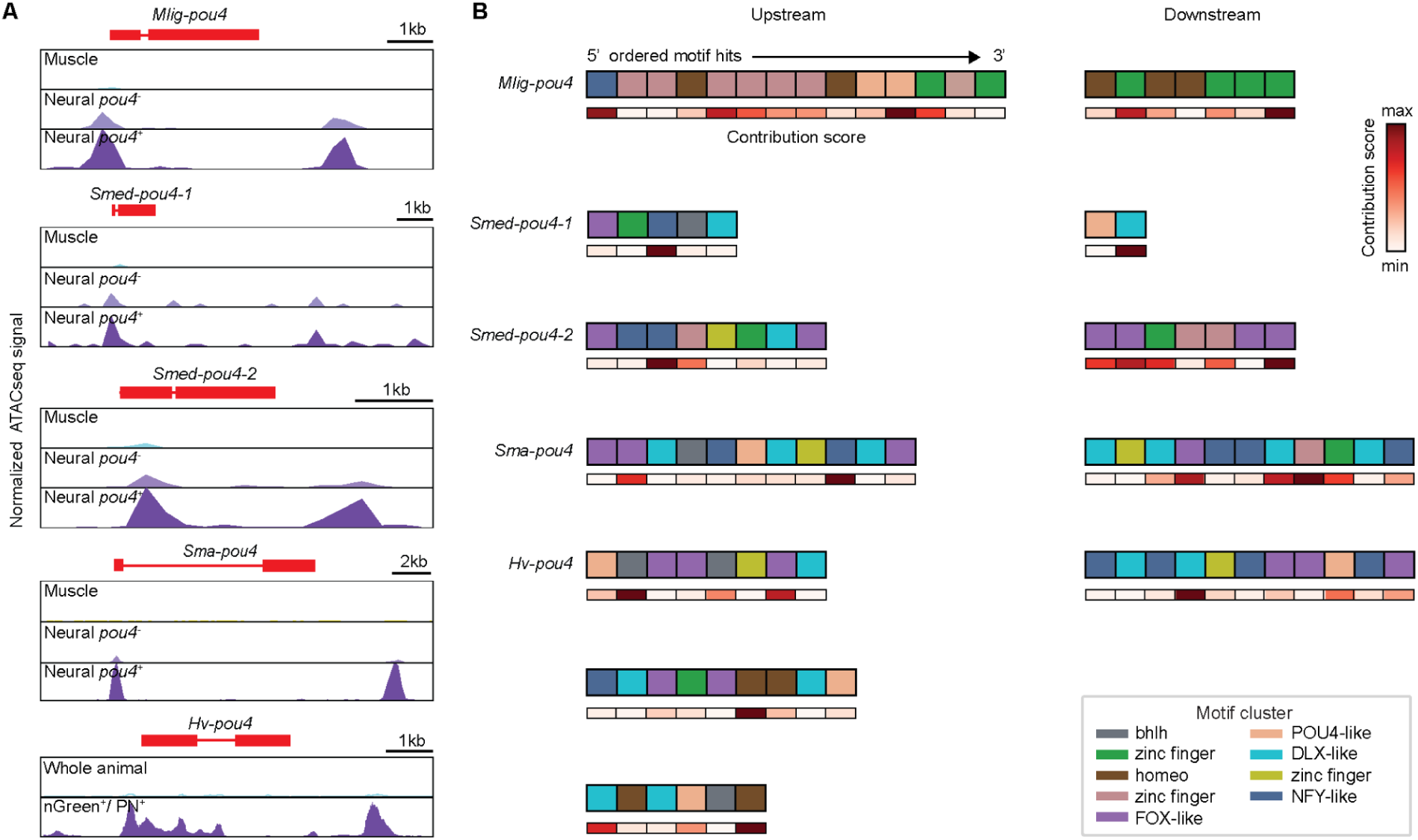
Extensive turnover of regulatory features associated with a deeply conserved transcription factor. (A) Genome tracks near the *pou4* loci for the three flatworms and *H. vulgaris*. *N. vectensis* data is too sparse at this locus and excluded in this analysis. Aggregated ATACseq coverage of *pou4*^+^ neural clusters are compared to *pou4*^-^ neurons and muscles for flatworms or the whole-animal sample in *Hydra*. *pou4* gene tracks are indicated in red, on the plus strand. (B) Representations of high-importance motif hits in the upstream (left) and downstream (right) peaks associated with *pou4* genes in each species. Individual motif hits are represented as boxes in the order they appear in the peaks. Boxes are colored to indicate motif types, grouped by sequence similarity (see **Methods**). Below these, 1D heatmaps indicate the relative contribution of each motif determined by the neural models.

### Synonymous motifs specify accessibility biases towards cell type families

Our findings support a model where drawing motifs from a large ‘vocabulary’ with conserved cell type family biases is sufficient to maintain accessibility of a sequence within those families. While species-specific combinatorial motif usage may define detailed cell type-level accessibility, family-level accessibility should be independent of this. Since large-scale transgenic experiments are impractical in most non-model species, we tested this hypothesis *in-silico* by using ChromBPNet models trained on one species to predict the accessibility of peak sequences from another species. Specifically, we compared *S. mansoni* and *S. mediterranea* models, as their genomes exhibit comparable GC content, reflected by their similar Tn5 biases (**Figure S3A**). These models were fed with peak sequences from the other species to predict accessibility and underlying contribution scores (**Figure 5A**), allowing us to compare family-level accessibility patterns and the motifs driving them.

**Figure 5.**
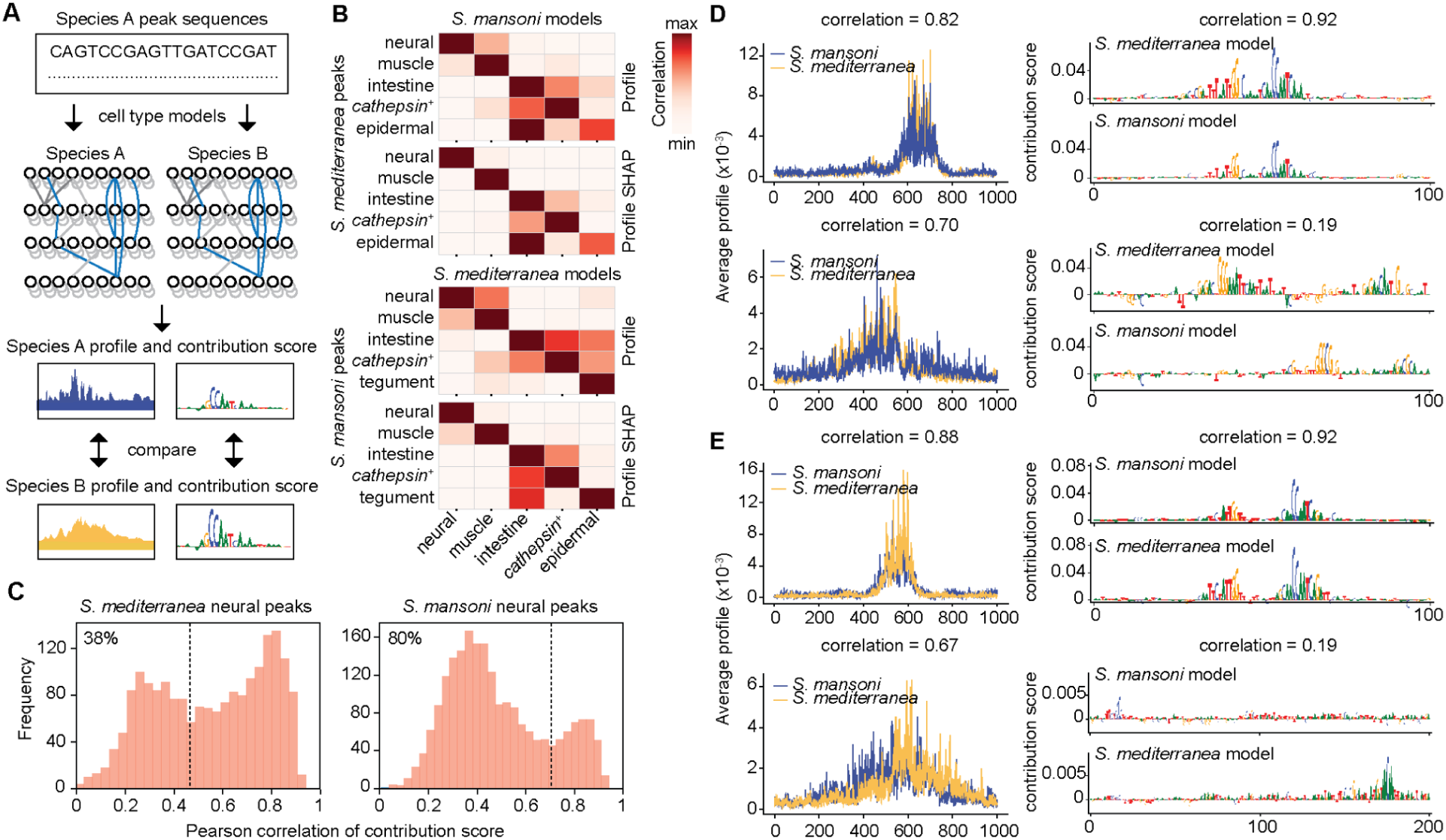
Cross species model predictions suggest synonymous motifs are sufficient to convey cell type family biases. (A) Schematic outlining cross-species model predictions in which models trained on one species are used to predict and interpret the accessibility of peak sequences from another. These outputs are compared with models trained on data from the species of origin. (B) Heatmaps showing the performance of cross-species model prediction on family marker peaks, with *S. mansoni* models predicting *S. mediterranea* peaks (top), and *S. mediterranea* models predicting *S. mansoni* peaks (bottom), as measured by the relative strengths in correlations of predicted profiles and contribution scores (see **Methods**). (C) Histograms showing bimodally distributed SHAP score correlations for peaks with highly correlated accessibility predictions between species (>0.55). Dashed lines separate the populations with high and low SHAP correlations, showing 38% of *S. mansoni* model predictions and 80% of *S. mediterranea* model predictions below these thresholds. (D-E) Predicted accessibility profiles (left) and model contribution scores (right) are compared between species for example peaks with highly correlated accessibility predictions and either highly (top) or lowly (bottom) correlated SHAP scores when testing the *S. mansoni* neural model on *S. mediterranea* sequences (**D**), or the *S. mediterranea* neural model on *S. mansoni* sequences (**E**).

Strikingly, we found that the accessibility predictions made by models trained on matching families were most strongly correlated across species (**Figure 5B**). We hypothesized that, while these predictions may rely on similar sets of motifs overall, the exact sequences recognized by the model to predict accessibility of a given peak should be more flexible between species. To test this, we compiled neural-specific peaks with similar accessibility predictions from neural models of both species (accessibility correlation > 0.55) and computed the correlation of their respective SHAP scores (**Figure 5C**). We found the correlations were bimodally distributed, suggesting that, although the family-biased accessibility of these sequences remained recognizable across species, large fractions (∼40%-80%) of predictions were made based on different underlying sequence features, or motifs, per peak (**Figures 5C-E**). This result implies that these motifs convey synonymous meanings in specifying genome accessibility across species, in support of our model.

## Discussion

In this work, we demonstrated that genome regions accessible in specific cell type families use motif sets, or vocabulary, conserved over long evolutionary distances, and that the usage of these motifs reveals conserved hierarchical relationships between cell types. However, the combinatorial use of these motifs in individual cell types, which show clear patterns within individual species, is almost entirely non-conserved. Even master regulators with highly conserved expression and function (e.g., *pou4*) can have distinct sequence features driving local accessibility.

Our findings suggest that regulatory sequences may encode multiple levels of information, balancing stability and specificity. While combinations of motifs can drive cell type-specific accessibility patterns, mutations may disrupt these complex sequences rapidly^9,10^. In contrast, we propose a model of ‘collective maintenance’: maintaining accessibility by drawing motifs from a broad ‘vocabulary’ is more robust to mutations but can only convey simpler information, such as the division of cell type families (**Figure 1B**). This layered information allows sequences to evolve within family-level constraints while exploring novel cell type-specific patterns through combinatorial motif use. Our comparisons of distantly related species allowed us to identify these highly stable evolutionary constraints.

This mode of conservation could emerge through opportunistic TF binding. In regions under evolutionary pressure to remain accessible in particular cell type families, following mutations disrupting a TF binding site, the accessibility can be stabilized by transient interactions with other DNA binding proteins expressed within that family. These interactions may then become reinforced as further mutations accumulate. However, this local turnover does not account for the observed strong, global preferences for certain motif combinations that differ between species. The combinatorial usage may be shaped by TF-TF interactions that are highly context-dependent^52^, evolve faster than DNA-binding specificity, and may bias the set of DNA binding partners available to form such stabilizing interactions in a species-dependent manner. Understanding how protein-DNA and protein-protein interactions collectively determine the genome accessibility landscape remains an important area for future research.

Beyond protein-DNA interactions, other factors can also modulate genome accessibility, such as chromatin remodelers, non-coding RNAs, and chemical modifications that alter DNA occupancy and chromatin states^53^. Indeed, many of our WC motifs may reflect the influences of these factors. The core of the ‘collective maintenance’ concept is that genome access is maintained by factors drawn from a large, conserved pool, with selective pressure on maintaining access, rather than the specific factors that achieve it, and thus this principle may be generalizable to various types of regulatory factors.

## Acknowledgements

*S. mansoni* (strain: NMRI) was provided by the NIAID Schistosomiasis Resource Center for distribution through BEI Resources, NIH-NIAID Contract HHSN272201000005I. We thank Eugene Berezikov, Jakub Wudarski, and Lisa Glazenburg for sharing the *M. lignano* NL12 strain, the experimental protocols for animal husbandry, and the genome assembly before publication, Jochen Rink and Andrei Rozanski for sharing the planarian genome assembly and annotation, Celina Juliano and Abby Primack for sharing the *Hydra* ATACseq data before publication. We thank Kevin Wang, Detlev Arendt, Eileen Furlong, Jacob Musser, and William Greenleaf for stimulating discussions, Mohamed Ameen, Souradeep Sarkar, and Livia Wyss for experimental help, Laksshman Sundaram, Shulin Mao, Ruohan Ren, and Hunter King for help with the analysis, Steve Granick, Liqun Luo, and Celina Juliano for critical reading of the manuscript, and all Wang group members for feedback. C.C. is supported by a Stanford DARE fellowship. This work is supported by an NIH grant (1R35GM138061), the Neuro-omics project of Wu Tsai Big Ideas in Neuroscience program, a Stanford Bio-X Interdisciplinary Initiatives Seed Grant (IIP R11-40), and a HFSP grant (RGY0085/2019).

## Author contributions

CC, JG, and BW designed the research. CC, PL, and JG performed the experiments. CC and JG analyzed the data. APampari, APatel, and AK provided advice on ChromBPNet models. CC, JG and BW wrote the paper with input from all other authors. AK and BW acquired the funding. AK and BW supervised the project.

## Data and code availability

Single-cell multiome and ATACseq datasets generated in this study are available in the Sequence Read Archive (SRA) and can be accessed using BioProject number, PRJNA1153627. All code used in this study can be found on GitHub at https://github.com/Bo-Wang-Lab/cell_type_regulatory_evolution. This same code base, alongside all of the processed data and ChromBPNet models necessary to reproduce our analyses will be available via Dryad.

## Methods

### Animal care and maintenance

*M. lignano* strain NL12 were cultured in artificial sea water (ASW) (37 g/L MarineMix Pro - Bulk Reef Supply, Cat# 208269) with diatom *Nitzschia curvilineata*, at 20 °C and 60% humidity, in a 14 hr/10 hr light/dark cycle^54^. The diatom was seeded in 150 mm glass dishes, and grown in F/2 media (Bigelow, Cat# MKf250L) for approximately one week, until confluent. The F/2 media was poured off, and replaced with fresh ASW into which animals were transferred once per week. The animals were starved in ASW for 48 hr prior to sample preparation for sequencing experiments.

Sexual and asexual *S. mediterranea* were maintained in 0.75× Montjuic salts or 0.5 g/L Instant Ocean Sea Salts (IO) supplemented with 0.1 g/L sodium bicarbonate, respectively, in the dark at 18 °C, and fed calf liver paste weekly. The animals were starved for 7 days prior to sequencing experiments.

Juvenile and adult *S. mansoni* were collected from infected Swiss Webster mice (NR-21963) 3.5 weeks and 7 weeks post infection, respectively, by hepatic portal vein perfusion using 37 °C DMEM supplemented with 5% heat-inactivated FBS^19^. Worms were cultured in Basch Media 169 supplemented with 1× Antibiotic-Antimycotic at 37 °C, 5% CO_2_. All experiments with and care of mice were performed in accordance with protocols approved by the Institutional Animal Care and Use Committees (IACUC) of Stanford University (protocol approval number 30366).

### Nuclei isolation

Approximately 70 *M. lignano* whole animals, aged between 4-6 weeks, or 150 heads or tails 3 hours post amputation (hpa) were transferred to a 2 mL tissue grinder (Fisher Scientific, 50-212-708) and washed with artificial seawater. Heads and tails were collected by relaxing the animals in 1:1 7.14% MgCl_2_ (Fisher BP214-500):ASW and cutting tissues between male and female gonads with a No. 15 scalpel 3 hr before nuclei isolation. Animals were dounced 30 times with pestle B (Fisher Scientific, 50-212-708) in 1 mL lysis buffer (10 mM Tris-HCl, 10 mM NaCl, 3 mM MgCl_2_, 0.1% Tween-20, 0.1% NP-40, 0.01% digitonin, 1% BSA, 1 mM DTT, supplemented with 1 U/μL RNase inhibitor (Sigma-Aldrich, 3335402001)), and filtered through a 40 μm cell strainer. Samples were incubated for 10 min, pipetting up and down 10 times every 2 min. 9 mL wash buffer (10 mM Tris-HCl, 10 mM NaCl, 3 mM MgCl_2_, 1% BSA, 0.1% Tween-20, 1 mM DTT, 1 U/μL RNase inhibitor) was then added to the nuclei suspension, and the sample was centrifuged at 600× g for 5 min at 4 °C. The nuclei were resuspended in 60 μL of diluted nuclei buffer (10x Genomics) supplemented with 1 mM DTT and 1 U/μL RNase-inhibitor, and filtered through a Flowmi 40 μm filter (Sigma-Aldrich, BAH136800040) before moving to library prep.

Five mid-sized (∼5-7 mm in length) asexual planarians were amputated into three pieces using a No. 22 scalpel either 6 hr before or immediately prior to nuclei isolation. Animals were rinsed in IO three times and transferred to a 7 mL Dounce-All-Glass tissue grinder (DWK Life Sciences, 885300-0007). The animals were dounced in a 1 mL lysis buffer using pestle B (DWK Life Sciences, 885300-0007) continuously for 3 min (∼48 strokes) on ice and further incubated for 7 min. Remaining tissue chunks were removed through serial filtration using 100 μm, 70 μm, and 40 μm cell strainers. The filtered suspension was distributed into 100 μL aliquots, each of which were diluted 1:10 with wash buffer, and centrifuged at 800× g for 5 min at 4 °C. The nuclei were then resuspended in 1 mL wash buffer and centrifuged again at 800× g for 5 min. The nuclei were finally resuspended in 100 μL diluted nuclei buffer, supplemented with 1 mM DTT and 1 U/μL RNase-inhibitor, and filtered through a 30 μm cell strainer before moving to library prep.

Tissue fragments surrounding the testes were dissected from five mid-sized (∼7-10 mm in length) sexual planarians to enrich for GSCs and reproductive cell types. The fragments were rinsed three times in 0.75× Montjuic salts and transferred to a 7 mL Dounce-All-Glass tissue grinder. The tissues were dounced 5 times with pestle A and 17 times with pestle B. The tissues were further broken down by pipetting 10-15 times before passing through a 40 μm cell strainer. The suspension was incubated on ice for 10 min, pipetting up and down 5 times every minute, and washed in 9 mL wash buffer, and centrifuged at 350× g for 5 min at 4 °C. Nuclei were resuspended in 2 mL wash buffer, passed through 100 μm, 70 μm and 40 μm filters sequentially, then centrifuged at 350× g for 5 min at 4 °C. The nuclei were finally resuspended in 80 μL of diluted nuclei buffer, supplemented with 1 mM DTT and 1 U/μL RNase-inhibitor, and filtered through a 30 μm cell strainer before moving to library prep.

Thirty adult schistosomes were chopped into multiple pieces in Basch Media either 6 hr before or immediately prior to nuclei isolation. These adult tissues or 60 uninjured juvenile parasites were washed three times in DPBS (Cytiva, SH30028.03) and placed in a 7 mL Dounce-All-Glass tissue grinder with 1 mL of nuclei lysis buffer. The animals were dounced with pestle B continuously for 3 min (∼48 strokes), and further lysed on ice for 7 min. The homogenized tissue was then passed through a 30 μm filter to remove undissociated tissue chunks. The nuclei suspension was distributed in 100 μL aliquots, diluted with 900 μL of wash buffer, and centrifuged at 800× g for 5 min at 4 °C. The nuclei were then resuspended in a diluted nuclei buffer before moving to library prep.

### Library preparation and sequencing

An aliquot of the isolated nuclei was stained with propidium iodide (1:500, 1 mg/mL stock) to inspect nuclei quality and confirm complete lysis before library preparation. Nuclei from adult *S. mansoni* parasites were processed using the 10x Genomics Chromium Controller and Chromium single-cell V1 ATAC-seq kit. Amplified libraries were quantified using bioanalyzer, and sequenced using Illumina HiSeq 4000 (150 bp paired-end), with coverages around 27,000 read pairs per nucleus. Nuclei from *M. lignano*, *S. mediterranea* and juvenile *S. mansoni* were processed using the 10x Genomics NextGEM Single Cell Multiome ATAC + Gene Expression Kit. Amplified libraries were quantified using bioanalyzer and sequenced using Illumina NovaSeq6000, with coverages ranging from 20,000-30,000 read pairs per nucleus for ATACseq libraries and 15,000-35,000 read pairs per nucleus for RNAseq libraries. Mean coverages for individual libraries are listed in **Table S1**.

### RNAi

Gene fragments for preparing dsRNA were amplified from cDNA using oligonucleotide primers listed in **Table S2** and cloned into the vector pJC53.2 (Addgene Plasmid ID: 26536), followed by in vitro transcription as previously described^55^. 10-15 juvenile parasites were soaked in Basch 169 media supplemented with ∼20 μg/mL dsRNA for 2 weeks^19^. The media containing dsRNA were refreshed daily. In all RNAi experiments, dsRNA of the ccdB and camR insert sequence in the pJC53.2 plasmid was used as the negative control. Each RNAi was performed at least twice with two technical replicates each time to assess phenotype.

After treatment, juvenile parasites were pulsed with 10 μM of EdU (TCI Chemicals) overnight, and killed in 6 M MgCl_2_ for 30 s, fixed with 4% formaldehyde supplemented with 0.2% Triton X-100 and 1% NP-40 for 4 hr, and then dehydrated in methanol. Dehydrated animals were bleached in 3% H_2_O_2_ in methanol for 30 min, rehydrated with 50% methanol in PBSTx (0.3% Triton-X in PBS), followed by two PBSTx washes. Samples were permeabilized by 10 μg/mL proteinase K for 20 min, and then post fixed with 4% formaldehyde. EdU incorporation was detected by click chemistry reaction with Carboxyrhodamine 110 Azide conjugates (Click Chemistry Tools).

For fluorescence imaging, samples were mounted in scale solution (30% glycerol, 0.1% Triton X-100, 2 mg/mL sodium ascorbate, 4 M urea in PBS) and imaged on a Zeiss LSM 800 confocal microscope using a 40× water immersion objective (LD C-Apochromat Corr M27). Fluorescence images are maximal intensity orthogonal projections, representative of all images taken in each condition.

### Multiome data processing and quality control

Raw ATACseq reads from all three species were mapped to their respective genome assembly (Mlig_4_5 for *M. lignano*^56^, dd_Smes_g4 for *S. mediterranea*^57^, and SM_V7 for *S. mansoni*^58^) using Chromap^59^ with default parameters and ‘–preset atac –drop-repetitive-reads 10’. The output fragment files were then read into ArchR version Release_1.0.1^60^ as arrow files through createArrowFiles. At this stage, gene activity score matrices and genome-wide tiled accessibility matrices were computed by setting addGeneScoreMat=TRUE and addTileMat=TRUE. Likely doublets were identified and removed using addDoubletScores and filterDoublets. Low-quality nuclei were further filtered by thresholding the total number of fragments per nuclei and transcription state site (TSS) enrichment according to the parameters in (**Table S1**).

The raw cell-by-gene RNAseq counts matrix for *M. lignano* was generated using kallisto-bustools^61^ (kb count with -x 10XV3) mapping to the Mlig_4_5 v5 reference transcriptome^56^. Raw reads from *S. mediterranea* and *S. mansoni* were aligned to dd_Smes_g4, and SM_V7 respectively using STARsolo^62^. RNA counts were added to the ArchR projects using addGeneExpressionMatrix, which included a sum normalization to 10,000 reads per nucleus (CP10k). Iterative Latent Semantic Indexing (LSI)^63^ was performed independently on the gene expression matrix and genome tile matrix for dimensionality reduction to 30 dimensions for each modality using addIterativeLSI. These reduced dimension embeddings were concatenated through the function addCombinedDims (except for *S. mansoni* adult data which used the ATAC representation only). Harmony^64^ was applied to the resulting joint embeddings using addHarmony, in order to reduce batch effects.

We clustered nuclei in the batch-corrected joint embedding using Seurat’s^65^ implementation of Louvain clustering^66^. Some clusters were later merged based on shared SAMap mappings between species (see below). For *S. mansoni* adult cells, RNA data from Wendt et al.^16^ was integrated using addGeneIntegrationMatrix. Clusters were annotated using both gene activity scores and gene expression of known cell type markers (**Figure S1A**). Finally, we generated a 2D representation using uniform manifold approximation and projection (UMAP).

### Peak calling and annotation

Pseudo-bulk fragment files for each major cluster were generated using FilterCells from Signac^67^ before peak calling with MACS2 using ENCODE ATACseq pipeline^68^. Peaks called for each cluster were merged using an iterative overlap method^69^ to produce a consensus peak set for each species. The consensus peaks were annotated using ChIPseeker^70^ with default parameters, except tssRegion = c(-1000,1000) and genomeAnnotationPriority = c("Promoter", "Exon", "Intron", "3UTR", "5UTR","Downstream", "Intergenic").

Cell-by-peak accessibility matrices were exported from ArchR and further processed using PeakVI^71^. PeakVI models were trained with a sample identifier set as batch_key, and all other parameters left as default. The models’ latent representations of the datasets were exported to be used in metacell identification (see below) for *M. lignano* and *S. mansoni*. PeakVI’s differential_accessibility method was run with default parameters to identify putative marker peaks for each cluster. Differentially-accessible peaks were filtered by thresholding prob_da at least 0.85 and effect_size at least 0.1.

### Metacell analysis

For *S. mediterranea*, the CP10k-normalized gene expression matrix was exported from ArchR and further processed through scVI^72^. An scVI model was trained with a sample identifier set as batch_key, and all other parameters left as default. This model’s latent representation of the dataset was exported to be used for metacell identification.

Metacells were computed separately for each sample using SEACells^73^, operating on the reduced dimension embeddings from PeakVI for *M. lignano* and *S. mansoni*, or from scVI for *S. mediterranea*. One SEACell was identified for every 75 nuclei in the *M. lignano* and *S. mansoni* data, and for every 50 nuclei in the *S. mediterranea* data. Individual nuclei whose maximum SEACell assignment score was less than 0.1 were discarded before aggregating ATACseq and RNAseq reads among nuclei assigned to the same metacell. Metacells were assigned cluster annotations according to the majority annotation of their member nuclei.

### Testing the predictive power of feature sets

Logistic regression classifiers were trained to predict one-vs.-rest cluster labels given per-cell or per-metacell measurements of peak accessibility or motif deviations, respectively. Models were constructed through scikit-learn’s^74^ LogisticRegression class with parameters penalty=’l2’, C=1e5. Predictive power of the feature set for that label was then measured using the Matthews Correlation between the true and predicted labels in a held-out test set. For each cluster, this process was repeated for 10 train-test splits, with 70% of the observations being used for training and 30% for testing.

### SAMap

CP10k-normalized gene activity score matrices were exported from ArchR and processed using the SAM algorithm^75^ with Harmony batch correction enabled. For *M. lignano*, only data collected from uninjured animals were used for mapping to avoid longer runtimes with the larger dataset. For *S. mediterranea*, nearest neighbor-averaged gene scores were used after applying the ImputeMatrix function in ArchR to compensate for lower ATACseq coverage. Processed SAM objects from each species were used to run SAMap^6^, and ArchR clusters that were indistinguishable by their mapping scores between species were merged to produce the final cluster annotations. For mapping between cell type families, clusters were grouped based on family annotations, and these family labels were compared through SAMap, with alignment scores less than 0.3 left out for plotting. Gene scores of neurons or muscles were processed similarly and mapped separately based on subcluster labels, plotting scores above 0.2.

### ChromBPNet models

We trained ChromBPNet models using a pre-release version (v1.3-pre-release) from the ChromBPNet Github repository^76^. Pseudo-bulk fragment files for each tissue were generated as described above, and adjusted to have +4/-4 shifts required by the ChromBPNet pipeline. Per-base coverage tracks of 5’ read insertions were then generated in BedGraph format using bedtools^77^ and converted into BigWig format with bedGraphToBigWig. To ensure sufficient coverage depth for rare cell types (∼5 million reads), we merged the following coverage tracks: for *M. lignano*, protonephridial and anchor cells; for *S. mediterranea*, pharynx, protonephridia, *ophis^+^* and GSC progeny; for *S. mansoni*, oesophageal gland cells were merged with protonephridia, and S1 with vitellocytes.

We first used the build_pwm_from_bigwig.py script to estimate Tn5 bias motifs for each species from the ATACseq coverage. We then trained Tn5 bias models to allow ChromBPNet to regress out these biases during final training. A single bias model was trained for both *S. mediterranea* and *S. mansoni* datasets using reads from the schistosome muscles, as these species share similar genomic GC content and Tn5 bias motifs (**Figure S3A**). A separate bias model was trained for the *M. lignano* dataset using reads from its muscles, as the higher GC content resulted in a distinct bias motif (**Figure S3A**). Muscle data were chosen in both cases for its high read depth. The Tn5 bias model architecture followed the ChromBPNet defaults except for stride=100. These models were trained on low-coverage, non-peak regions in order to ensure the bias was only learned at closed regions to represent noise-based cut sites. We ran TF-MoDISco-lite on the trained models to verify that no known TF binding motifs were learned. These learned motifs were later passed into final trained ChromBPNet models (described below) to confirm proper accounting of biases, where maximum profile response should be <0.003.

We trained ChromBPNet models for each major tissue type in all species using their respective peak and GC-matched non-peak sequences as input. All ChromBPNet models were trained with default parameters, except for the *S. mansoni* models, which used a filter length of 9 bp for the first input convolutional layer to improve model performance. The performance of ChromBPNet models were evaluated through correlation between observed ATACseq insertions and ChromBPNet predicted counts, as well as profile prediction accuracy using Jensen-Shannon distance (JSD) between measured ATACseq profiles and ChromBPNet predicted profiles. To test the stability of all our models, we trained two additional models for each tissue type using separate train-test splits over genome contigs (three-fold validation), and found that the performance metrics were comparable across folds (**Figures S3A** and **S5D**). For downstream analysis, we selected a fold in which the models across tissue types were most optimal. All training was performed using Nvidia A100 GPUs with CUDA v11.0.

We applied DeepSHAP implementation of DeepLIFT^78^ to the trained ChromBPNet models to estimate the predictive contribution of each base in a peak sequence to the predicted total counts and profile signals. TF-MoDISco-lite v2.0.0^36^ was then applied to identify regions with high counts contribution across consensus peaks with the default parameter except final_min_cluster_size=10, trim_to_window_size=15, and max_seqlets_per_metacluster= 200,000. High-contribution sequences were clustered based on within-group contributions and sequence similarity and consolidated into position frequency matrices (PFMs).

### Motif annotation

Pairwise distances were calculated for all PFMs output from MoDISco. These distances were defined as the sum of the per-position JSD for the aligned PFM pair with the offset and orientation (forward or reverse complement) which minimizes that sum. Similarity scores were then calculated as the correlations between the rows of the pairwise distance matrix. Motifs were clustered hierarchically using a complete linkage of the similarity matrix, and clustering was terminated when merging two clusters proposed by this linkage would result in an average cluster PFM that has an average per-position JSD > 0.2 from one or more member PFMs. These clustered PFMs were mapped to the JASPAR2020 non-redundant core motif database using TomTom^79^, and clusters whose consensus sequence exactly matched a motif in the database were removed from the analysis. Specifically, 152 learned motifs exactly matched the PFMs in JASPAR and were removed to avoid redundancy. The same clustering procedure was applied to the JASPAR motifs themselves, and the combined set of PFM clusters were used for peak annotation. The motif clusters were named with a prefix of either JC (for those derived from JASPAR PFMs) or WC (for those derived from MODISCO), followed by a unique 4 digit number, the name of the most similar JASPAR motif, and the total JSD value associated with that mapping. We only considered JSD<0.5 as reliable matches.

Motif hits in peak sequences were annotated using ArchR’s addMotifAnnotation with default parameters except cutOff=1e-5, and average contribution scores within each hit were computed for each ChromBPNet model. To understand how these values compared with the models’ responses to non-meaningful sequences, the sequences were shuffled within each peak while maintaining their dinucleotide frequency distributions. These shuffled sequences were then interpreted by each model. Average contribution scores within the regions annotated as motif hits were calculated as above to produce a background distribution per model. To identify motif hits that strongly influenced observed accessibilities, we selected instances with contributions above the 97th percentile of the background distribution for *M. lignano*, 95th percentile for *S. mediterranea*, and 98th percentile for *S. mansoni* under at least one model’s interpretation.

To combine motif hits around *pou4* into similarity classes (**Figure 4B**), we normalized the contribution scores per peak to have unit root-mean-squared (RMS) values, and extracted weighted one-hot encodings of each motif hit. Similarities between each hit pair were then calculated as the convolution between their weighted encodings with maximum value over all possible shifts and orientations (i.e., forward vs. reverse complement). The hits were then hierarchically clustered based on a complete linkage of the pairwise correlations between rows of the similarity matrix, and the resulting tree was manually trimmed to produce groups of sequences with high similarity and annotated based on representative motif hits in TomTom^79^.

### Marginal footprinting

Consensus sequences for motifs of interest were inserted into the center of randomized 2114-bp non-peak regions to create synthetic sequences. Profile probability predictions were generated for both the forward and reverse complements, and then averaged to obtain a footprint for each synthetic sequence. To attain a marginal footprint, we generated predictions for 256 synthetic sequences and averaged the predictions. A broad accessibility peak with a dip at the motif inserted region is characteristic of TF binding^28^. For these *in-silico* footprint analyses, we inserted several tandem motif sequences as individual motifs were generally insufficient to induce clear responses.

### Motif deviations and marker motif identification

chromVAR deviations were calculated using a local python implementation of the method described by Schep et al.^38^. Deviations were calculated based on the SEACell-aggregated ATACseq reads for each dataset. All deviation values used in these analyses were deviation z-scores, calculated over 50 GC-matched background peak sets identified in ArchR using the addBgdPeaks method.

Marker motifs were identified using a one-sided Mann-Whitney U test to compare chromVAR deviations between SEACells belonging to each cluster against all others. P-values derived from these tests were adjusted to account for multiple testing bias using the Benjamini-Hochberg method. A motif was considered to be a marker for a given cluster if the adjusted p-value for the associated test was less than 0.1, and the median deviation value within the cluster was greater than 1. For cell type family markers, we combined motif markers of at least one member of the family. The average accessibility of the motif in that family was then computed as the median deviation value of all SEACells belonging to member clusters for which that motif is a valid marker.

### Co-occurrence analysis

For each pair of motifs, we calculated the number of peaks in which both motifs are annotated. In this process, multiple hits to the same motif were not considered co-occurring if they overlapped within a peak, and similarly, hits to distinct motifs that overlapped by 3 or more bases were not considered to co-occur. We then define enrichment as the ratio of the fraction of peaks containing a sequence hit to both motifs divided by the product of the fractions of peaks containing sequence hits for each motif. To avoid false positives between rare motifs, any pairs with an expected and observed number of peak hits less than or equal to 2 were ignored and assigned an enrichment value of 1, corresponding to expectations based on random chance.

### Subcluster analysis

Nuclei belonging to neural clusters were re-processed using PeakVI for *M. lignano* and *S. mansoni*, and scVI for *S. mediterranea*. The resulting latent representations of the neural data were used to run Leiden clustering^80^. First, a k-nearest-neighbor (knn) graph was computed on this embedding using scanpy’s^81^ implementation of bbknn^82^ (sc.external.pp.bknn) with a sample identifier set as batch_key, neighbors_within_batch=10, and trim=0. Leiden clustering was then performed using scanpy’s sc.tl.leiden method. The resolution parameter for clustering was adjusted independently for each dataset until the resulting clusters were all predictable with Matthews Correlation above ∼0.8 using logistic regression on the ATACseq counts (for *M. lignano* and *S. mansoni*) or sum-normalized and log2-transformed RNAseq counts (for *S. mediterranea*). This gave resolutions of 0.13 for *M. lignano*, 0.2 for *S. mediterranea*, and 0.3 for *S. mansoni*. All other clustering parameters were left as default. These subcluster labels were then propagated to the SEACells data by majority vote. Muscle clusters were processed similarly, with resolution parameters of 0.3 for *M. lignano*, 0.8 for *S. mediterranea*, and 0.45 for *S. mansoni* used in the final Leiden clustering.

### Motif module analysis

Marker motifs were identified for each neural subcluster. Highly variable motifs were defined as those with a standard deviation of chromVAR deviations specifically in the neurons greater than 1 for all three flatworm species. Highly variable motifs that were also markers of at least one neural subcluster in all three species were retained for module analysis. The Mann-Whitney test results were used to create a binarized subcluster-by-motif accessibility matrix for each dataset. Pairwise Jaccard similarities were computed for the motifs per species and modules were identified as connected components of the graph with edges between motif pairs with identical binary accessibility patterns (Jaccard similarity = 1). Non-trivial modules containing more than one motif are depicted in **Figures 3B** and **S4E**, and a complete list of module assignments is available in **Table S3**.

### Cnidarian data analysis

*H. vulgaris* raw data were obtained from Primack et al.^47^ and aligned to AEP genome assembly^83^. *N. vectensis* raw data were obtained from Sebe-Pedros et al.^48^ and aligned to Nvec200^84^. Raw sequencing reads were filtered for sequencing adaptors and low-quality reads using Trimmomatic^85^, and then mapped to the corresponding genome using Bowtie2^86^. Ambiguously mapped reads were filtered using samtools view (with -u -F 524 -f 2), followed by filtration of PCR duplicates using Picard MarkDuplicates with default parameters^87^.

Aligned reads were then shifted by +4/-4 bp for training ChromBPNet models. For *H. vulgaris* data, we combined four replicates of Tg(actin1:GFP)^rs3-in^ ATACseq libraries with two replicates of Tg(tba1c:mNeonGreen)^cj1-gt^ libraries as the neural population, which we termed, PN^+^/nGreen^+^, and three replicates of AEP1 libraries as the whole animal control. For *N. vectensis* data, we merged two replicates of *elav^+^* ATACseq libraries as the neural population, and two replicates of *elav^-^* libraries as the non-neural population. Finally, separate bias models were trained for both the *H. vulgaris* and *N. vectensis* datasets using their whole-body or non-neural controls despite the similarity in their estimated Tn5 bias motifs in order to compensate for differences in the experimental techniques used to collect these datasets.

Motifs annotation, filtration and deviation calculations were done as described above with a few changes. When filtering motifs, we selected motif hits with observed average contribution scores that were above 95th percentile for *H. vulgaris*, and 90th percentile for *N. vectensis* of the background distributions. When identifying marker motifs, a motif is considered to be enriched in the neural populations if the average deviation of a motif in the neural populations is greater than 1 and less than 0 in the whole animal control for *H. vulgaris* and non-neural population for *N. vectensis*, respectively. To identify differentially accessible peaks, we used DiffBind^88^ to calculate normalized read counts across all peaks. UROPA^89^ was then employed to annotate all ATACseq peaks based on nearest TSS. EdgeR^90^ was then used to find differentially accessible peaks between conditions.

### Gene phylogeny

In addition to *M. lignano*, *S. mediterranea*, and *S. mansoni*, homologs of the planarian *pou4* gene (SMESG000030759.1) were identified in reference transcriptomes of *Schistosoma japonicum* (HuSJv2^91^), *Macrostomum hysterix* (Machtx_SR1_v2^92^), *Macrostomum cliftonense* (Maccli_GV23d_v2^92^), *Schmidtea polychroa* (dd_Spol_v4^93^), *Prostheceraeus crozieri* (PRJEB44148^94^), *Stenostomum brevipharyngium* (go_Sbre_v1^95^), *Pristina leidyi*^96^, *Platynereis dumerilii* (PdumBase v2^97^), *Convolutriloba longifissura*^98^, *Hofstenia miamia* (HmiaM1^99^), *Nematostella vectensis* (Nemve1^100^), and *Hydra vulgaris* (HVAEP^83^) combining results from OrthoFinder^101^ (run with default parameters) and genes sharing EggNOG annotations^102^. The protein sequences of these genes were aligned using Clustal Omega^103^ with default parameters. A tree model was fit to the alignment using IQ-TREE^104^ with the model finder plus option using parameters -seed 0 -bb 1000 -alrt 1000 -m MFP+G -mset raxml. The optimal model found based on the Bayesian Information Criterion (BIC) was JTT+F+R4. This gene tree was then adjusted to reconcile with the known species tree using GeneRax^105^ with --strategy SPR. Finally, the resulting trees were visualized using phylo.io^106^. This same procedure was applied to homologs of the planarian *etv4* (SMESG000008885.1) and *pax6-like* (SMESG000004009.1) genes, with JTT+F+R4 and VT+F+R5 models, respectively.

## Supplemental figures

**Figure S1.**
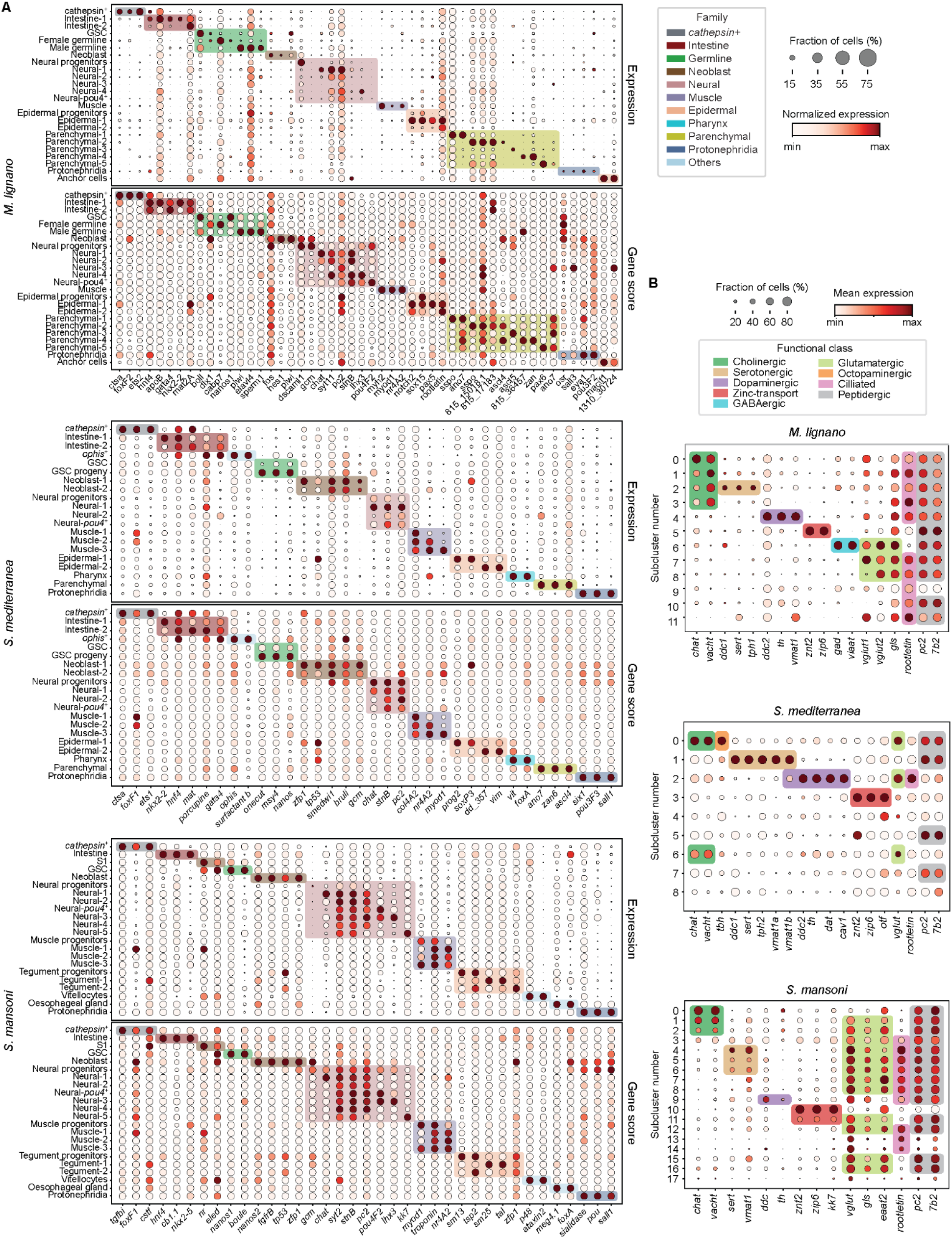
Marker genes used in cell type and functional class annotation, related to Figures 1 and 3. (A) Dot plots showing min-max normalized average expression (top) and gene activity scores (bottom) of selected marker genes used in annotating clusters. *Macrostomum* parenchymal populations (sharing *ano7* and *ascl4* expression with the planarian) express known markers of secretory glands such as the prostate and cement gland^22^. Clusters are grouped into cell type families highlighted with colored blocks. Contig numbers for all mentioned genes are listed in Table S2. (B) Dot plots showing min-max normalized average expression of functional neural markers in neural populations for *S. mediterranea* and *S. mansoni*. For *M. lignano*, min-max normalized average accessibility of peaks near each of these genes is shown instead. Functional classes of these subclusters are indicated by colored boxes.

**Figure S2.**
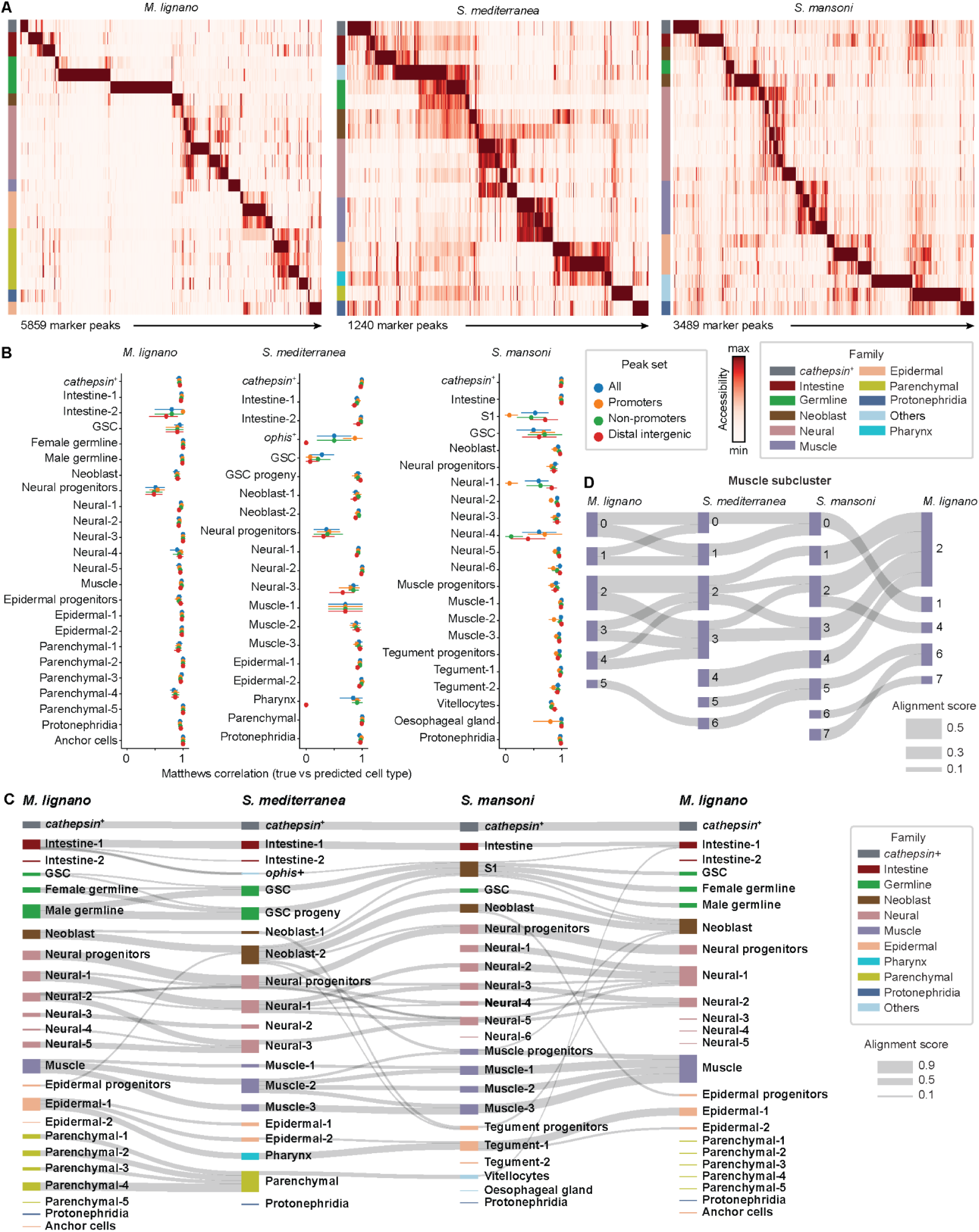
Loss of cell type specificity in conserved accessibility patterns, related to Figure 1. (A) Min-max normalized average accessibility values for the top 5% most specific marker peaks of each cluster, colored by their cell type family on the left. (B) Matthew’s correlation of logistic regression models trained separately on promoter, non-promoter (including exonic, intronic, and intergenic peaks), or distal, intergenic peaks alone in predicting cluster identities, showing no systematic difference in predictive power between these peak sets. (C) Sankey plot summarizing the SAMap mappings between all clusters based on gene scores, with blocks representing clusters colored according to their families. Mappings are degenerate but are largely self-contained within families, while clusters containing species-specific cell types remain unconnected. (D) Sankey plot summarizing the SAMap mappings between flatworm muscle cell types. Muscle types separate into three major shared classes as well as a fourth which appears to have been lost in *S. mediterranea*. Mappings between types within each class are degenerate.

**Figure S3.**
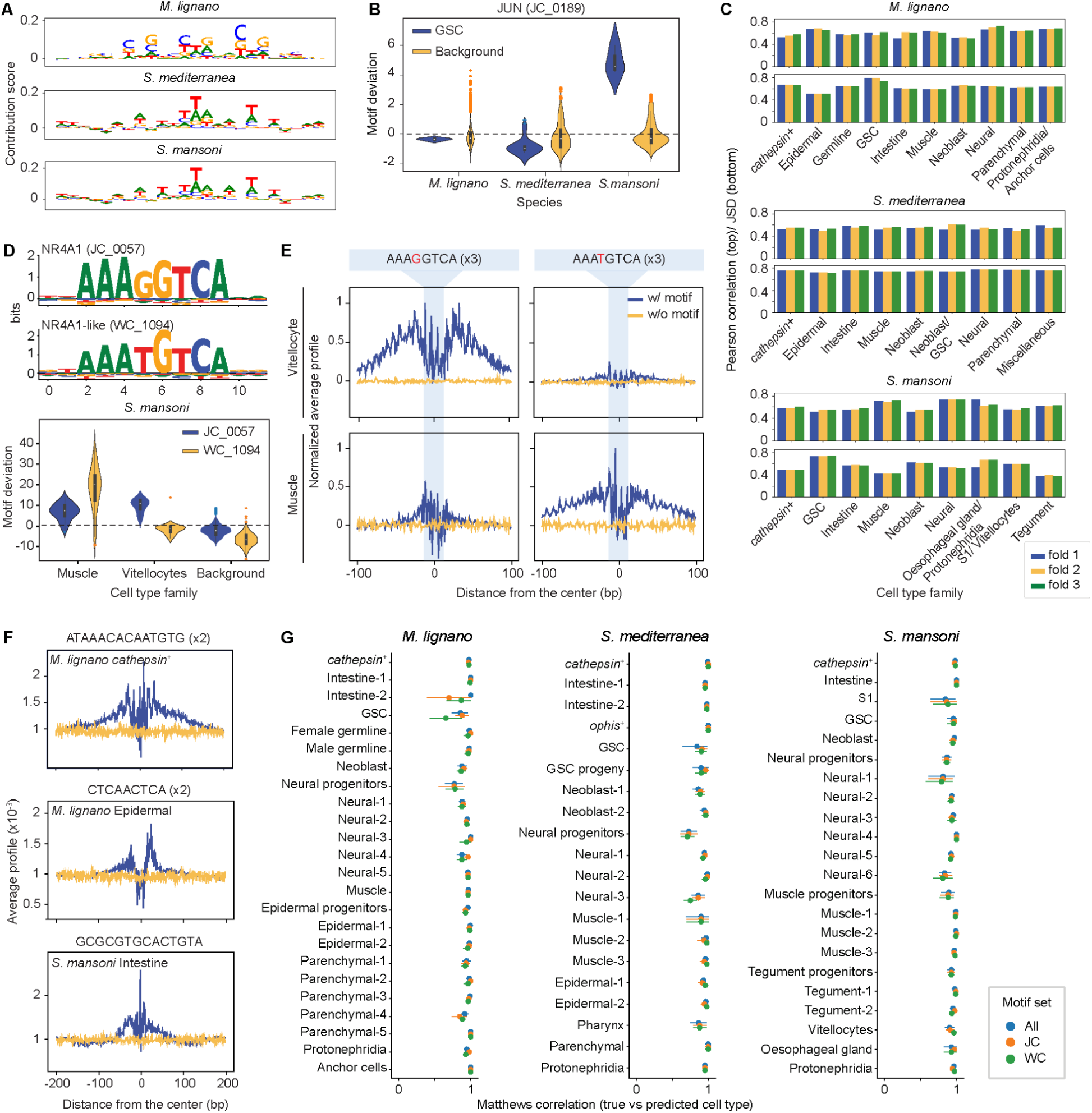
ChromBPNet models learn the sequence features dictating ATACseq signal, related to Figure 2. (A) Representative Tn5 motif logos extracted from *M. lignano*, *S. mediterranea*, and *S. mansoni* data. (B) Violin plots showing motif deviations of JUN motifs (JC_0159) in the accessible genome regions of GSC metacells (blue) compared with all other cells (yellow) across the three flatworms. Note that the GSC enrichment is specific to the schistosome. (C) Pearson correlation between true and predicted insertion counts (top) and JSD between observed and predicted ATAC insertion profiles at base resolution across all peaks in held out contigs over 3 train-test splits (3-fold validation) for all trained ChromBPNet models. (D) Violin plots showing motif deviations in metacells (bottom) for an NR4A1 motif derived from the JASPAR2020 database (JC_0057) and a similar motif with a single base substitution (WC_1094) learned from the schistosome muscle model. WC_1094 is highly accessible in muscles specifically, while JC_0057 has lower accessibility overall and is expanded to the vitellocytes. (E) *In-silico* footprinting using the schistosome muscle (bottom) and vitellocyte (top) models predicting the accessibility of a tandem (3×) consensus sequence for JC_0057 (left) or WC_1094 (right) inserted at the center of 256 randomized 2114-bp background sequences, showing the average prediction over these sequences. Both models show a footprint-like response to the JC_0057 sequence, albeit lower for the muscle model, whereas the vitellocyte model does not respond to the WC_1094 sequence. The footprint analysis results are consistent with motif deviation patterns in D. We insert tandem motifs because single motifs are generally insufficient to induce clear responses. (F) Footprints of example WC motifs listed in **Figure 2D** by the models from which they are learned. *In-silico* footprinting performed as in E using tandem (2×) consensus sequences. Footprint-like responses suggest these sequences may interact with DNA binding proteins. (G) Matthew’s correlation of logistic regression models trained separately on JC or WC motifs in predicting cluster identities, showing no difference between these two motif sets.

**Figure S4.**
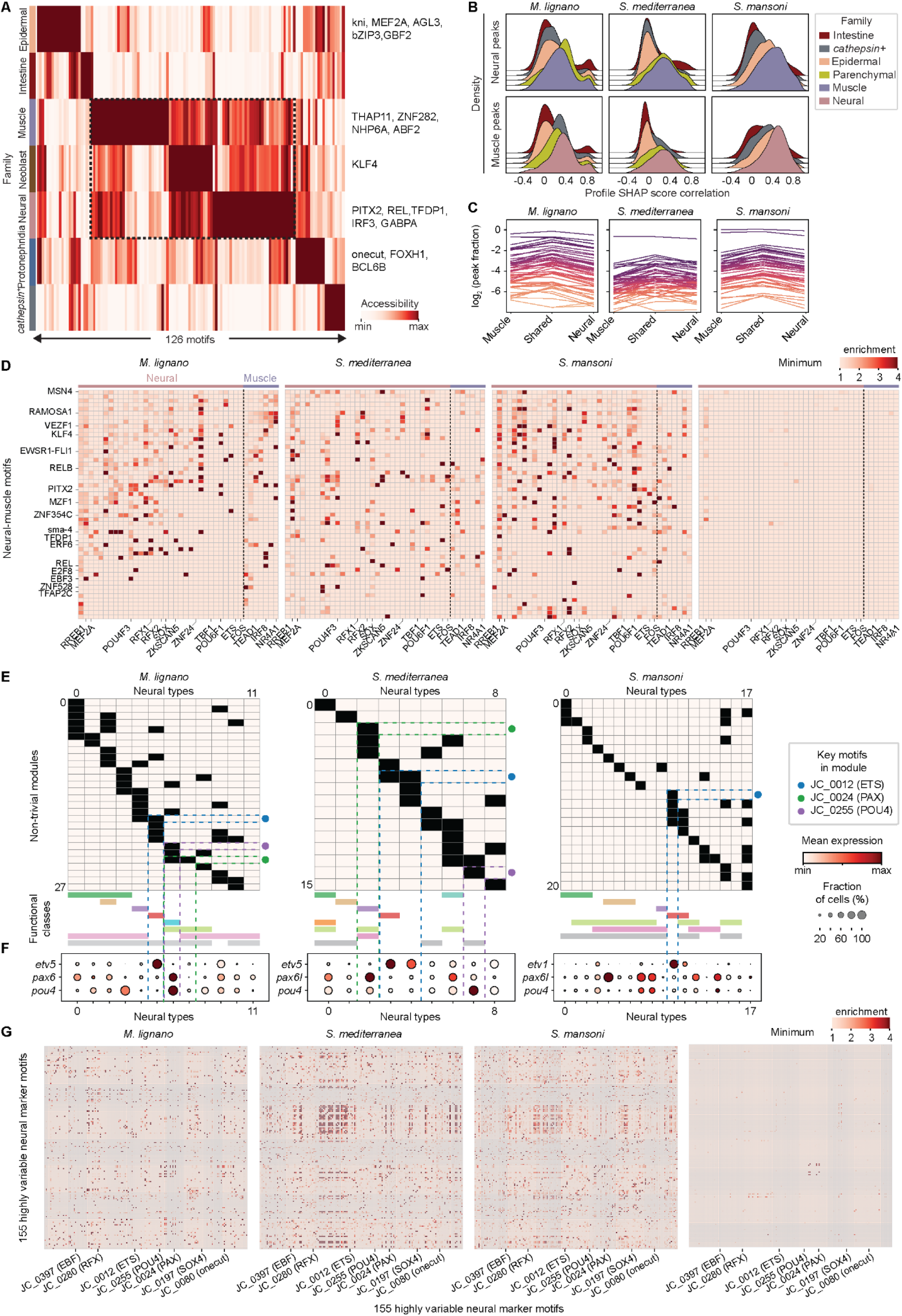
Patterns of motif use in neurons, muscles and parenchymal cells, related to Figure 3. (A) Min-max normalized average motif deviations for 126 parenchymal marker motifs (columns) shared between *M. lignano* and *S. mediterranea* (**Table S3**) in 7 cell type families (rows) in the *S. mansoni* data. Example motifs with high accessibility in each family are indicated to the right. Note the bias towards neurons, muscles, and their progenitors (dashed box). (B) Top: histograms showing the correlations between SHAP scores from different models predicting accessibility of neural marker peaks with the SHAP scores from neural models for each species. Bottom: corresponding plots for muscle marker peaks. Note that the muscle and neural models always have the best cross-family model predictions for each other’s peaks. (C) Fractions of family-specific and neural-muscle shared marker peaks containing hits to motifs shared by neurons and muscles in each species. Individual motifs are represented as a matched set of 3 connected points. These motifs are clearly present in family-specific markers, despite a slight increase in abundance within shared marker peaks. (D) Heatmaps showing the enrichment of co-occurrence between motifs shared by neurons and muscles and neural- and muscle-specific motifs. Motifs are ordered by decreasing the number of sequence hits within the *M. lignano* genome from top to bottom and left to right. Minimum enrichment values across all three species are shown in the rightmost plot. (E) Binarized accessibility profiles of non-trivial motif modules (containing more than one motif) in the neural types of each species. Filled boxes indicate that motifs in a module (rows) are highly accessible in a given neural type (columns). Functional classes for each type are labeled with colored bars at the bottom, and membership of several key motifs (ETS, PAX, and POU4) is indicated by colored dots to the right. (F) Dot plots showing min-max normalized average expression of TFs corresponding to the key motifs shown in E. *M. lignano* data is presented using the accessibility of peaks near each gene. (G) Pairwise co-occurrence plots for 155 highly variable neural marker motifs in all three species. Note the lack of conservation of motif-motif interactions between species, as indicated by the negligible minimum co-occurrence values in the rightmost plot.

**Figure S5.**
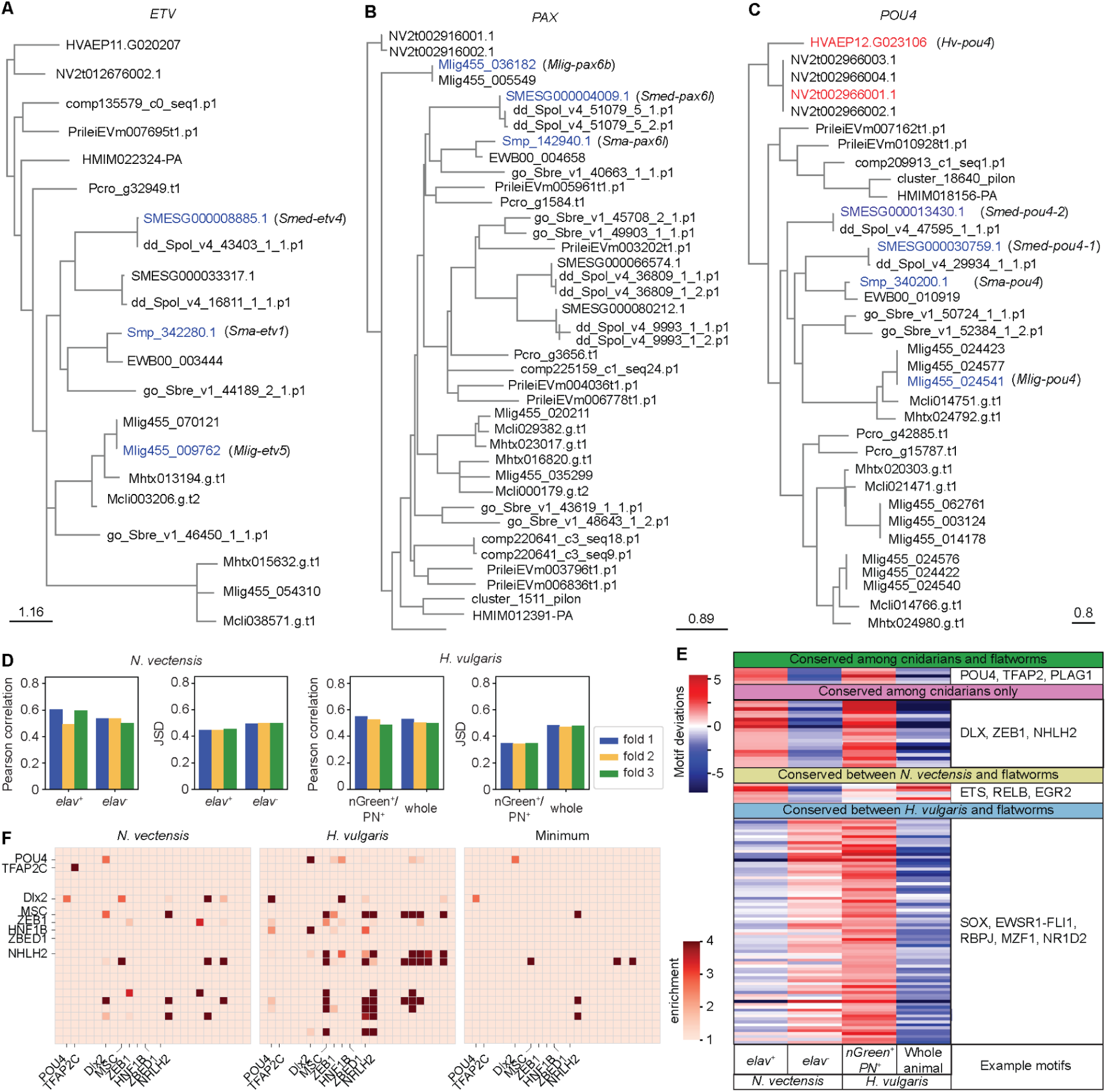
Comparison of neural TFs and motif use between flatworms and Cnidarians, related to Figures 3 and 4. (A-C) Phylogeny of *etv* (**A**), *pax* (**B**), and *pou4* (**C**) homologs in the three flatworm species with several intermediates (see **Methods**). Gene IDs for each species are prefixed as follows - SMESG: *S. mediterranea*, Smp: *S. mansoni*, Mlig455: *M. lignano*, dd_Spol_v4: *S. polychroa*, EWB: *S. japonicum*, Mcli: *M. cliftonense*, Mhtx: *M. hysterix*, Pcro: *P. crozieri*, go_Sbre_v1: *S*. *brevipharyngium*, Prilei: *P. leidyi*, comp: *Platynereis dumerilii*, cluster: *C. longifissura*, HMIM: H. miamia, HVAEP: H. vulgaris, NV2: N. vectensis. Flatworm genes included in **Figure S4** and **Figure 4B** are indicated in blue. Cnidarian genes included in **Figure 4B** are labeled in red. Scale bar: expected substitutions per site. (D) Pearson correlation between true and predicted counts (left) and JSD between observed and predicted profiles (right) in held out contigs over 3 train-test splits for *N. vectensis* and *H. vulgaris* models. (E) Heatmap showing average motif deviations of conserved neural motifs in neural (first and third columns) and non-neural (second and fourth columns) samples for *H. vulgaris*^47^ and *N. vectensis*^48^. Motifs shown are broken down into those shared by both cnidarians and all three flatworms (top first), those conserved between the cnidarians only (second), those shared by the flatworms and *N. vectensis* (third), and those shared by the flatworms and *H. vulgaris* (bottom). Example motifs in each group are listed to the right. (F) Co-occurrence of conserved neural motifs between the two cnidarians for each species, as well as the minimum co-occurrence between species.

## Supplemental Tables

Table S1. Sample condition, sequencing modality, sequencing depth, QC metrics for each sample.

Table S2. Information on key genes mentioned in this study.

(A) List of bZIP domain-containing genes in *S. mansoni* genome with primer sequences used for cloning in RNAi experiments.

(B) List of gene IDs associated with gene names used throughout the study.

Table S3. Information on key motif sets discussed in this study. See **Methods** for motif naming conventions.

(A) List of conserved motifs shared among all three flatworms for each cell type family. The smallest normalized accessibility values (min-max normalized over cell type families) across species are included for each motif in the family for which they are markers.

(B) List of parenchymal marker motifs shared between *M. lignano* and *S. mediterranea* and their accessibility in *S. mansoni* cell type families.

(C) List of motifs with conserved hierarchical use in neural-muscle superfamily. Module numbers are included for neural subcluster-defining motifs, corresponding to numbers shown in Fig. 3B and Fig. S4E for members of non-trivial modules.

(D) List of conserved neural motifs shared among cnidarians and flatworms, cnidarians only, between *N. vectensis* and flatworms, and *H. vulgaris* and flatworms.

## Supplementary Files

File S1. Numerical position frequency matrices (PFMs) of all motifs used in this study. PFMs are given a unique identifier prefixed with JC for those derived from JASPAR motifs alone, and prefixed with WC for those derived from MODISCO. Each PFM is annotated with the name of its most similar unclustered JASPAR motif, with the total per-position JSD for that alignment provided in parentheses.

File S2. Contribution score tracks for upstream and downstream peaks of *pou4* loci in *M. lignano*, *S. mediterranea*, *S. mansoni*, and *H. vulgaris* genomes. Tracks are provided as BigWig files which can be viewed online using the IGV app alongside the reference genome sequences and fasta indices that will be available via Dryad.

